# Single-cell analysis of localized low- and high-grade prostate cancers

**DOI:** 10.1101/2021.04.16.440238

**Authors:** Sebnem Ece Eksi, Alex Chitsazan, Zeynep Sayar, George V. Thomas, Andrew Fields, Ryan P. Kopp, Paul T. Spellman, Andrew Adey

## Abstract

Approximately, 30% of early-stage localized prostate cancer cases reoccur within 5 to 10 years [1, 2]. However, identifying precise molecular subtypes attributable to specific stages of prostate cancer has proven difficult due to high heterogeneity within localized tumors [3–5]. Bulk assays represent a population average, which is a result of the heterogeneity that exists at the individual prostate cancer cell level [6]. Here, we sequenced the accessible chromatin regions of 14,424 single-cells collected from 18 fresh-frozen prostate tumors using sci-ATAC-seq [7, 8]. We observed that shared chromatin features among low-grade prostate cancer epithelial cells were lost in high-grade tumors. Despite this loss, all high-grade tumors exhibited an enrichment for FOXA1, HOXB13 and CDX2 transcription factor binding sites within their accessible chromatin regions, indicating a shared trans-regulatory program. Single-cell analysis of the differentially accessible regions in high- versus low-grade prostate tumors identified two unique genes encoding neuronal adhesion molecules, NRXN1 and NLGN1. We found that NRXN1 and NLGN1 are expressed in the epithelial luminal, basal and neuroendocrine cells, as well as the immune, endothelial and neuronal cell types in all prostate tumors. Overall, these results provide a deeper understanding of the active gene regulatory networks in low- and high-grade prostate tumors at a striking resolution and provide critical insights for molecular stratification of the disease.

## INTRODUCTION

Tumor heterogeneity in prostate cancer poses a significant problem for molecular stratification of patients with localized prostate tumors [4, 9]. It is well established that only a subset of clinically identified prostate cancers lead to lethal metastatic disease [1, 10, 11]. However, significant heterogeneity within a specific tumor focus and across different tumor foci in the prostate gland results in complex evolutionary trajectories for the disease [3–5, 12]. The molecular heterogeneity within a tumor focus often leads to misclassification of the tumor grade and ineffective clinical treatment plans for many patients. The majority of the prostate cancer genomics and epigenomics data acquired to date originate from the bulk analysis of tumors that capture a population average of different cell types in the tumor [6, 13]. This generates a three-fold problem: 1) epithelial, endothelial, myeloid, lymphoid, nerve and other stromal cells that contribute to prostate cancer progression are reduced to a single component; 2) as a result, the dynamic bidirectional communication between these distinct components are not captured; 3) the heterogenous cell states within a single histopathological tumor grade are eliminated from the analysis.

Newly emergent single-cell technologies hold the key to profiling the vast heterogeneous landscapes of prostate cancer [8, 14–19]. Recently, whole genomes of 20 single cells from localized prostate tumors were sequenced [20], revealing significant cell-to-cell variation in mutations and complex subclonal trajectories. However, these microdissection-based studies do not sample enough cancer cells to represent an unbiased image of localized prostate tumors. Combinatorial indexing of single-cells provides a way to profile thousands of cells from various types of tissues [7, 21, 22]. Currently, this method has been applied to a select group of cancers [23, 24]. However, there are no current studies using these high-throughput single-cell technologies to characterize localized prostate adenocarcinoma.

To reveal the transformative changes in localized prostate tumors that lead to aggressive disease, it is important to capture the chromatin accessibility profiles of cells in low-grade and high-grade tumors. Open chromatin regions of cells contain not only promoter regions of actively transcribed genes, but also non-coding regulatory sequences. These sequences reflect the active gene regulatory networks that drive cell state transitions. Therefore, ATAC-seq (Assay for Transposase-Accessible Chromatin sequencing) technology provides a way to characterize both *cis*- and trans-regulators of cell states during tumor progression [25, 26].

The current best predictor of outcome in localized prostate cancer is the degree of differentiation, or grade of the tumor [27]. Grading of prostate cancer is reported through the Gleason score, which is a composite of the two most predominant Gleason grade patterns present in a sample [27]. Tumors that contain solely Gleason pattern 3 are often clinically indolent and tumors with higher Gleason pattern (≥4) are clinically significant and often associated with a much worse outcome. This association remains constant even if the higher pattern tumor foci make up a small proportion of the entire tumor population [27]. It should also be noted that if followed long enough, some of the patients that are diagnosed with Gleason pattern 3 tumors also develop aggressive disease under active surveillance [11, 28]. Therefore, determining the treatment strategies for patients with Gleason pattern 3 and 4 prostate tumors presents a clinical challenge [28–30].

Understanding the transition between indolent and aggressive disease requires determining the risk of progression [31]. Even though it is important to properly stratify tumors based on the Gleason pattern and score, confounding factors exist, such as the surgical margin status for patients who have gone through radical prostatectomy surgery, presence or absence of extracapsular extension and lymph node involvement. The Cancer of the Prostate Risk Assessment Post-Surgical (CAPRA-S) score and post-radical prostatectomy nomogram (Memorial Sloan Kettering Cancer Center) take these additional factors into consideration and provide ways to delineate the probability of disease recurrence [32–35].

Our primary objective in this study was to identify molecular and cellular markers associated with prostate tumors that have a primary Gleason pattern 3 (low-grade) and 4 (high-grade) to explore the heterogeneity of the localized disease. We used single-cell combinatorial indexing ATAC sequencing (sci-ATAC-seq) to characterize the *cis*- and trans-regulators of localized prostate tumors recovered from radical prostatectomies that contain Gleason pattern 3 and 4. We performed cyclic immunofluorescent (cyclic IF) microscopy to validate the expression of unique targets identified through sci-ATAC-seq at the protein level and to analyze the distribution of cell-types that express these targets in prostate cancer epithelial cells and in prostate tumor stroma.

## RESULTS

### sci-ATAC-seq identifies the accessible chromatin landscape of fresh-frozen prostate tumors at the single-cell level

We used combinatorial indexing to perform single-cell ATAC-seq (sci-ATAC-seq) from fresh-frozen prostate tumors collected from 18 patients via radical prostatectomy (Figure 1A). FFPE tissue blocks were prepared from each patient, and both H&E sections and unstained adjacent sections for immunofluorescent (IF) staining were obtained from each block (Figure 1A). Our cohort consisted of six low-risk, nine intermediate-risk and two high-risk patients based on their CAPRA-S scores and Memorial Sloan Kettering Cancer Center (MSKCC) nomograms (Supplementary File 1) [32, 34]. The majority of the tumors consisted of Gleason pattern 3 (low-grade) and 4 (high-grade) (Supplementary File 1).

**Figure 1:**
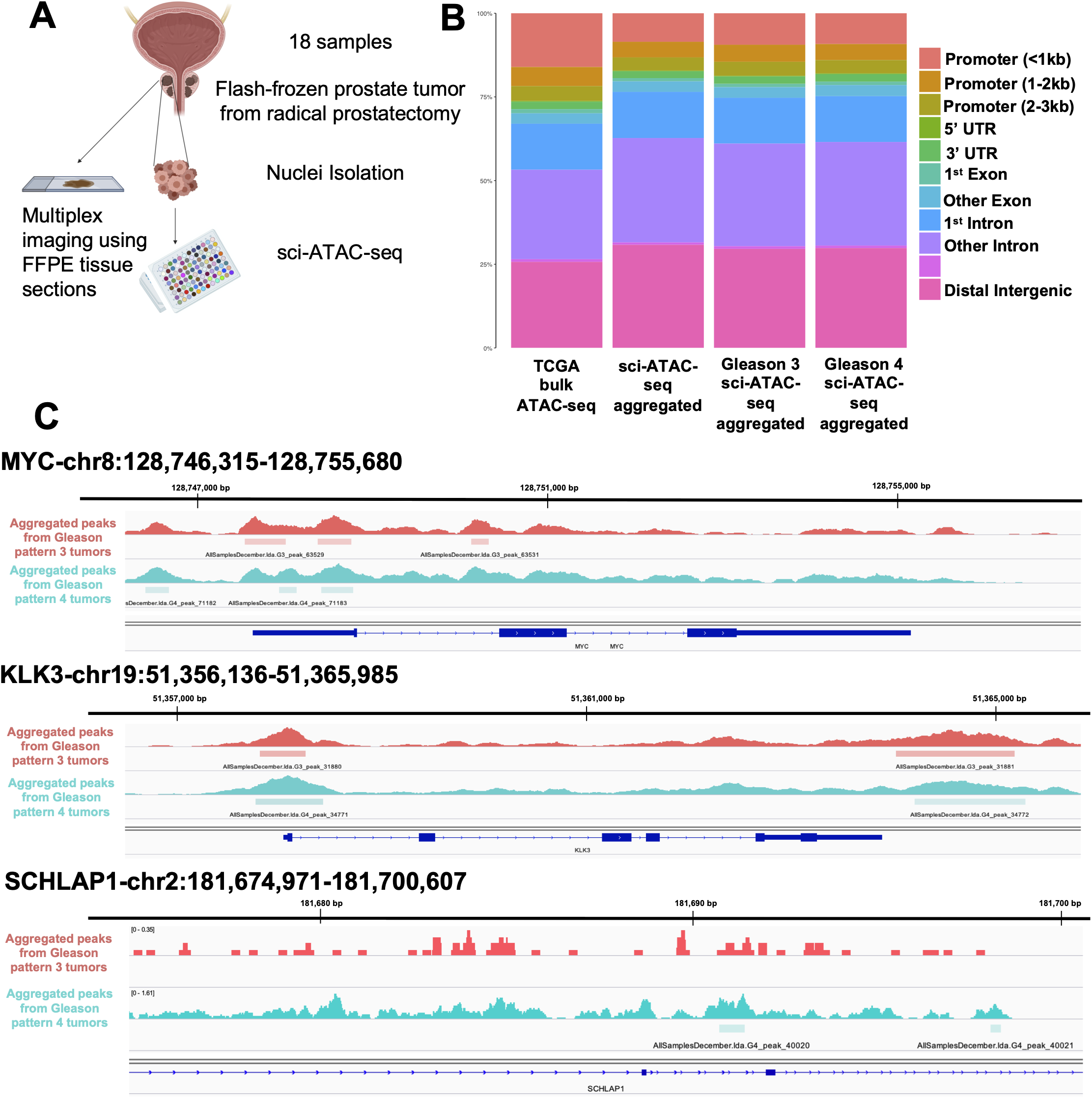
sci-ATAC-seq captures the chromatin accessibility landscape of prostate tumors with Gleason pattern 3 and 4. A. Diagram of experimental design. Flash-frozen prostate tumor samples were recovered from radical prostatectomies. Nuclei from each tumor were extracted and sorted into 96-well plates using unique combinatorial indices attached to transposase. FFPE tissue sections were collected from each patient along with an adjacent H&E section. B. Bar graph showing the percentage of peak distribution among functional genomic elements: promoter (<1kb, 1-2kb, 2-3kb), 3’UTR, 5’UTR, exon, intron, downstream (<300kb) and distal regions. All aggregated peaks from Gleason pattern 3 and 4 tumors and TCGA bulk ATAC-seq results are shown. C. sci-ATAC-seq peaks aggregated from Gleason pattern 3 (red) and 4 (blue) tumors. Called peaks are represented by dark blue rectangles. Examples of selected genomic regions: MYC gene (top), KLK3 (middle) and SCHLAP1 (bottom) are shown.

We analyzed the distribution of captured open chromatin regions across different functional genomic elements such as the promoter, 3’UTR, 5’UTR, exon, intron and distal regions (Figure 1B). We observed that aggregated sci-ATAC-seq data from all tumors exhibited a similar distribution across functional genomic elements when compared to bulk ATAC-seq results from the prostate adenocarcinoma (PRAD) TCGA data sets (Figure 1B, Supplementary Figure 1). This result indicated that sci-ATAC-seq was not biased in capturing open chromatin regions from specific genomic regions [36].

To confirm the quality of our acquired sci-ATAC-seq data sets, we aggregated peaks from Gleason pattern 3 and ≥4 tumors. We called 125,569 peaks and observed differentially accessible peaks around the promoter regions of several key genes in prostate cancer such as MYC and KLK3 (Figure 1C). To further validate the quality of our sci-ATAC-seq data sets, we examined differential accessibility to the region that encodes for SCHLAP1 long non-coding RNA in tumors with Gleason pattern ?4 as compared to pattern 3. SCHLAP1 is a prognostic marker that exhibits high expression in metastatic and high-grade prostate cancers [37, 38]. In agreement with this, our results showed higher accessibility to SCHLAP1 in Gleason pattern ≥4 tumors (Figure 1C).

Following several quality control procedures (Supplementary Figure 1, STAR Methods), 14,424 cells were recovered with high-quality sci-ATAC-seq reads from 18 primary prostate cancer samples (Figure 2A, Supplementary Figure 1). snapATAC analysis of our sci-ATAC-seq data, using the Latent Dirichlet Allocation method, identified 16 clusters of cells [39, 40]. Based upon cisTopic, each cluster of cells shared accessibility profiles grouped in 30 Topics (Figure 2B) [41]. 16 clusters were visualized by the Uniform Manifold Approximation and Projection (UMAP) (Figure 2A-B) [42, 43]. A heatmap of topic scores across clusters showed distinct chromatin accessibility profiles of cells (Figure 2C). Topics 4, 12 and 5 were mainly shared across cells from all clusters whereas Topics 20, 6 and 16 showed very specific profiles for a small number of clusters (Figure 2D). Additionally, we visualized putative gene transcription by quantifying the chromatin accessibility surrounding annotated transcription start sites (TSSs).

**Figure 2:**
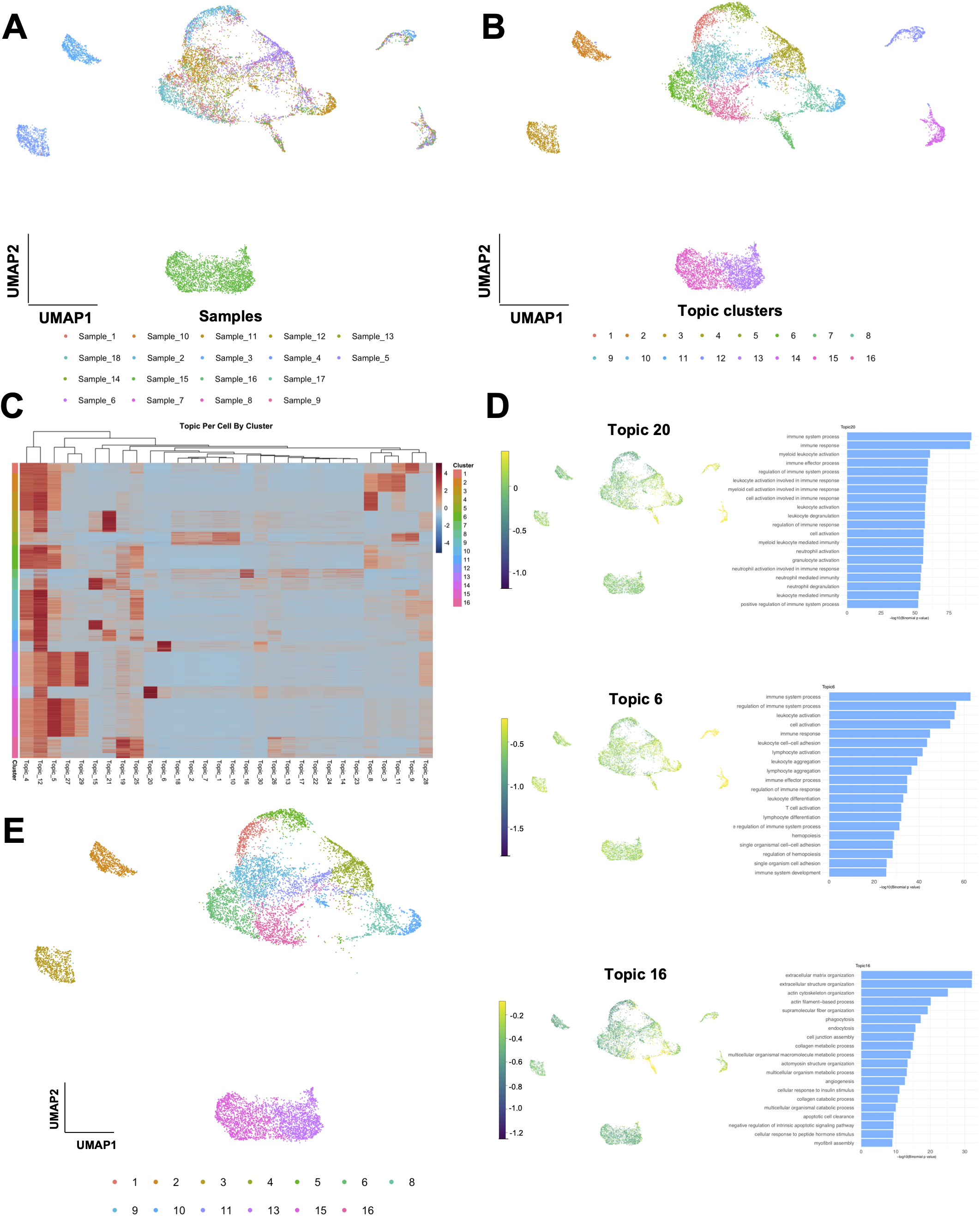
cisTOPIC identifiies 16 clusters of cells spanning 30 topics from 14,251 cells. A. Single-cells from 18 prostate samples were clustered using cisTOPIC based on their chromatin accessibility profiles. Cells are colored according to their sample IDs on the UMAP. B. Single-cells from 18 prostate samples formed 16 clusters based on their shared chromatin accessibility profiles. Cells are colored according to their cluster IDs on the UMAP. C. Heatmap showing the topic distribution across 16 clusters. D. UMAP showing Topic 20 was highly enriched in cells within Cluster 14 (top), Topic 6 was highly enriched in cells within Cluster 12 (middle) and Topic 16 was highly enriched in cells within Cluster 7 (bottom). GREAT analysis of GO profiles (right) for each Topic identified Cluster 14 as myeloid immune, Cluster 12 as lymphoid immune and Cluster 7 as stromal cells. E. Lymphoid and myeloid immune cells and stromal cells were removed from the UMAP for downstream analyses.

### Profiles from sci-ATAC-seq capture immune and stromal cell types in prostate tumors

Lymphoid, myeloid and other stromal cells have very different chromatin accessibility profiles compared to epithelial prostate cancer cells. Based on previous work, we expected that immune and stromal cell types associated with different prostate tumors would form distinct clusters of cells on the UMAP [44–46]. To examine this point, we used a cluster dendrogram of clusters to analyze the hierarchical relationship between the topics (Supplementary Figure 1). We observed Clusters 7, 12 and 14 attracted cells from all tumor samples that we isolated on the dendrogram (Figure 2A-B) and analyzed the topics enriched in each cluster. To identify the accessible chromatin regions that define these Clusters, we analyzed all Topics using Genomic Regions Enrichment of Annotations Tool (GREAT) that takes a set of genomic regions as its input, finds the associated *cis*-regulatory regions and outputs annotation terms that are significantly enriched within the input genomic regions [47]. Topics 20 and 6 showed an enrichment for GO terms related to the immune system process and immune response (Figure 2D). More specifically, immune Topic 20, which was enriched in cluster 14, contained GO terms associated with leukocyte and neutrophil activation, whereas immune Topic 6, which was enriched in cluster 12, contained GO terms associated with lymphocyte and T cell activation (Figure 2C-D). Similarly, we observed an enrichment for genomic regions associated with extracellular matrix organization for stromal Topic 16, which was enriched in cluster 7 (Figure 2D). Altogether, our results showed that cluster 7 consisted of stromal cells, cluster 12 consisted of lymphocytes and cluster 14 consisted of myeloid immune cells. We eliminated clusters 7, 12 and 14 from our downstream analyses to identify the gene regulatory network changes that specifically occur in epithelial prostate cancer cells (Figure 2E).

### Differentially accessible chromatin regions in single-cells are identified in Gleason pattern 4 prostate tumors

To delineate the chromatin accessibility profiles of epithelial prostate cancer cells, we examined the clustering of single cells captured from our patient cohort (Supplementary File 1). We observed that the majority of the cells extracted from low- and intermediate-risk prostate patients (Gleason 3+3, Gleason 3+4) formed a single large cluster in the dimensionally reduced UMAP space (Figure 3A). In contrast, single cells from one intermediate-risk (Gleason 4+4) and two high-risk (Gleason 4+4) prostate patients formed three distinct outer clusters (Figure 3A, Supplementary File 1). One intermediate-risk prostate tumor (Gleason 3+4), which consisted of 70% Gleason pattern 3 cells and 30% Gleason pattern 4 cells, had cells mostly clustering with low-grade prostate tumors, and a small group of cells clustering with high-grade prostate tumors (Figure 3A). Based on these results, we assigned single-cells to either Gleason pattern 3 or 4 categories on the UMAP (Figure 3B), to examine the *cis*-regulatory and trans-regulatory differences observed in cells from Gleason pattern 3 (low-grade, main cluster) and Gleason pattern 4 (high-grade, outer clusters).

**Figure 3:**
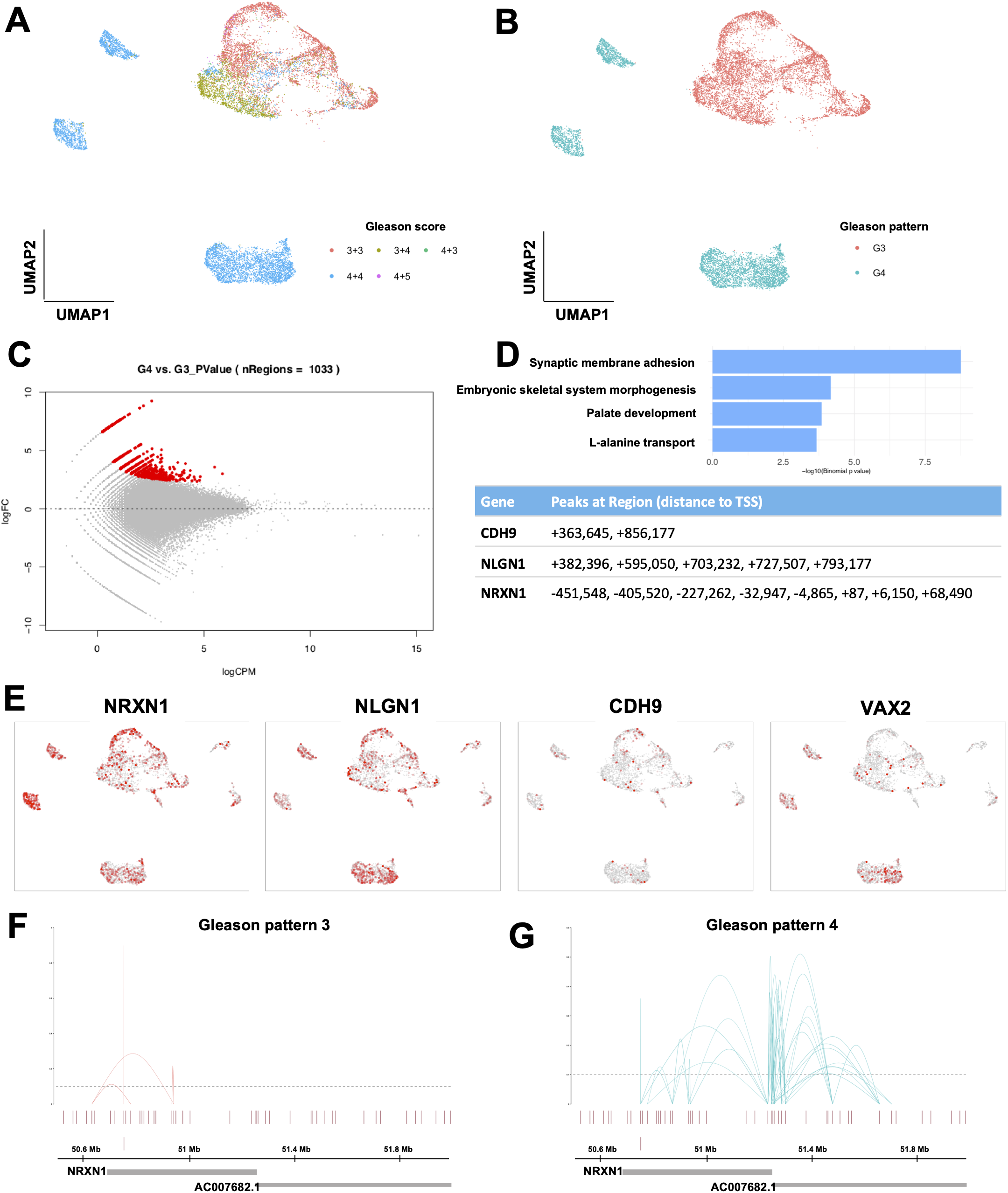
Tumors with Gleason pattern 4 have distinct chromatin accessibility profiles compared to tumors with Gleason pattern 3. A. cisTOPIC-UMAP of 18 prostate samples colored based on the Gleason score of each patient, which is a sum of two Gleason patterns: Gleason score 3+3 (red), Gleason score 3+4 (olive green), Gleason score 4+3 (green), Gleason score 4+4 (blue) and Gleason score 4+5 (magenta). B. Gleason pattern 3 single-cells are red, while 4 are blue and assigned according to their cluster. C. Distinct chromatin regions that are significantly more accessible in Gleason pattern 4 cells were identified (red). D. Gene ontology results of the differentially accessible regions are shown (top). 15 peaks were linked to three neuronal adhesion genes, NRXN1, NLGN1 and CDH9. (+) sign indicates 3’ distal sequences, (-) sign indicates 5’ distal sequences to the transcription start site (TSS). E. Examples of genes that had significantly more accessible chromatin regions in Gleason pattern 4 vs 3 cells. Each dot represents a single cell. Cells with accessible regions around the promoter of NRXN1, NLGN1, CDH9 and VAX2 are shown on the UMAP in red. F. The putative interactions between the distal regulatory regions and promoter sequences of NRXN1 in Gleason pattern 3 tumors (red). Co-accessibility scores are shown on the y-axis and the dotted lines represent the threshold. G. The putative interactions between the distal regulatory regions and promoter sequences of NRXN1 in Gleason pattern 4 tumors (blue). Co-accessibility scores are shown on the y-axis and the dotted lines represent the threshold. Cicero links around the NRXN1 loci were significantly higher in number in Gleason pattern 4 vs 3 tumors.

Differential accessibility functions of snapATAC (STAR Methods, [40]) identified top regions (Figure 3C, Supplementary Figure 2, Supplementary Figure 3) that are significantly accessible in Gleason pattern 4 tumors as compared to Gleason pattern 3 tumors. GREAT GO terms associated with accessible regions enriched in Gleason pattern 4 tumors included synaptic membrane adhesion, embryonic skeletal system morphogenesis, palate development and L-alanine transport (Figure 3D). Notably, genomic regions associated with the neuronal adhesion genes, NRXN1, NLGN1 and CDH9, were highly accessible in Gleason pattern 4 vs. 3 tumors (Figure 3D). It is important to note that UMAP shows accessibility to promoter sequences only, and does not show accessibility to the identified distal regulatory regions associated with these genes.

To determine if Gleason pattern 3 tumors would form separate clusters when analyzed in isolation, we eliminated all Gleason pattern 4 tumors from our analysis and performed topic analysis on Gleason pattern 3 tumors alone. We observed that Gleason pattern 3 tumors again formed one single cluster and did not form patient-specific clusters, suggesting certain chromatin accessibility features are shared across all Gleason pattern 3 tumors (Supplementary Figure 4).

We also profiled all cells based on their accessibility for known prostate cancer molecular subtypes (Supplementary Figure 5) [48]. Although patient specific patterns for ERG, ETV1, ETV4 and FLI1 were observed as expected, accessibility of these markers did not drive clustering of cells within the data (Supplementary Figure 5A). We demonstrated the accessibility profiles for some of the recurrently mutated genes MYC, SPOP, PTEN and IDH1 across all our samples (Supplementary Figure 5B). We observed higher accessibility to the promoter region of MYC in Gleason pattern 4 prostate tumors (Supplementary Figure 5B).

### Gleason pattern 4 prostate tumors are significantly enriched for neuronal adhesion molecules

Next, we used Cicero to examine the putative regulatory interactions around the NRXN1, NLGN1 and CDH9 loci [49]. Cicero is a sci-ATAC-seq method that finds putative regulatory interactions between regulatory sequences based on the coaccessibility of chromatin regions [49]. We observed an increase in the number of predicted *cis*-regulatory interactions around the NRXN1 locus in Gleason pattern 4 prostate tumors (Figure 3F). Several regions distal to the NRXN1 promoter and intronic sequences within the gene body that were inaccessible in Gleason pattern 3 tumors became accessible in Gleason pattern 4 tumors (Figure 3G). Even though not quantitative, this result was particularly striking since we had a smaller number of Gleason pattern 4 cells (5,334 cells) than Gleason pattern 3 (7,383 cells). We also observed some additional putative regulatory interactions in Gleason pattern 4 tumors around the NLGN1 and CDH9 loci, even though the increase in the number of links were not as high as around the NRXN1 locus (Supplementary Figure 5C).

To validate our results that show a significant enrichment in the chromatin accessibility profiles of high-grade tumors for neuronal adhesion molecules NRXN1, NLGN1 and CDH9, we analyzed the bulk ATAC-seq profiles using the TCGA PRAD data set [36]. TCGA PRAD data set consisted of 26 patients with intermediate- and high-grade prostate tumors (Supplementary Figure 6, Supplementary File 2). We observed a significant increase in the accessibility of NRXN1 and CDH9 chromatin sites in high-grade tumors as compared to intermediate-grade tumors (Supplementary Figure 6). Next, we analyzed the bulk RNA-seq profiles of 497 patients from the TCGA cohort and detected NRXN1 expression in the majority of the prostate tumors (Supplementary Figure 6). In contrast, we did not detect NLGN1 transcripts in the TCGA bulk RNA-seq data set (Supplementary Figure 6).

### Distinct trans-regulatory networks exist in Gleason pattern 3 and 4 tumors

sci-ATAC-seq gives a snapshot of all active genetic programs in individual cells [21, 50]. To understand the active gene regulatory networks in Gleason pattern 3 and 4 tumors, we aggregated all epithelial cells from Gleason pattern 3 and 4 tumors separately and analyzed the enriched TF binding motifs in each group using HOMER motif enrichment analysis [51]. We observed an enrichment for Fra1, Fra2, JunB, Atf3 and AP-1 in tumors with Gleason pattern 3 whereas tumors with Gleason pattern 4 showed an enrichment for FOXA1, HOXB13 and CDX2 (Figure 4A).

**Figure 4:**
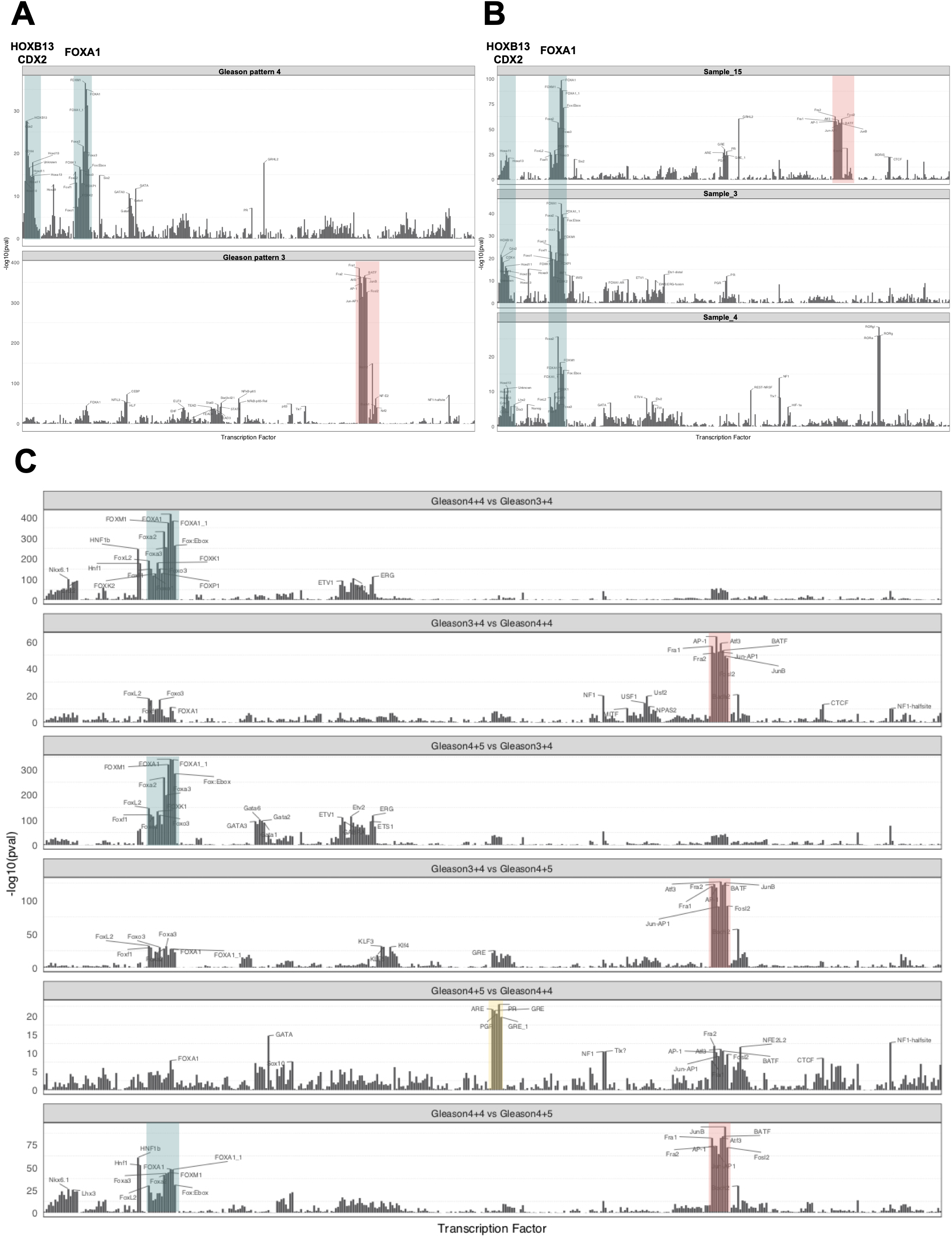
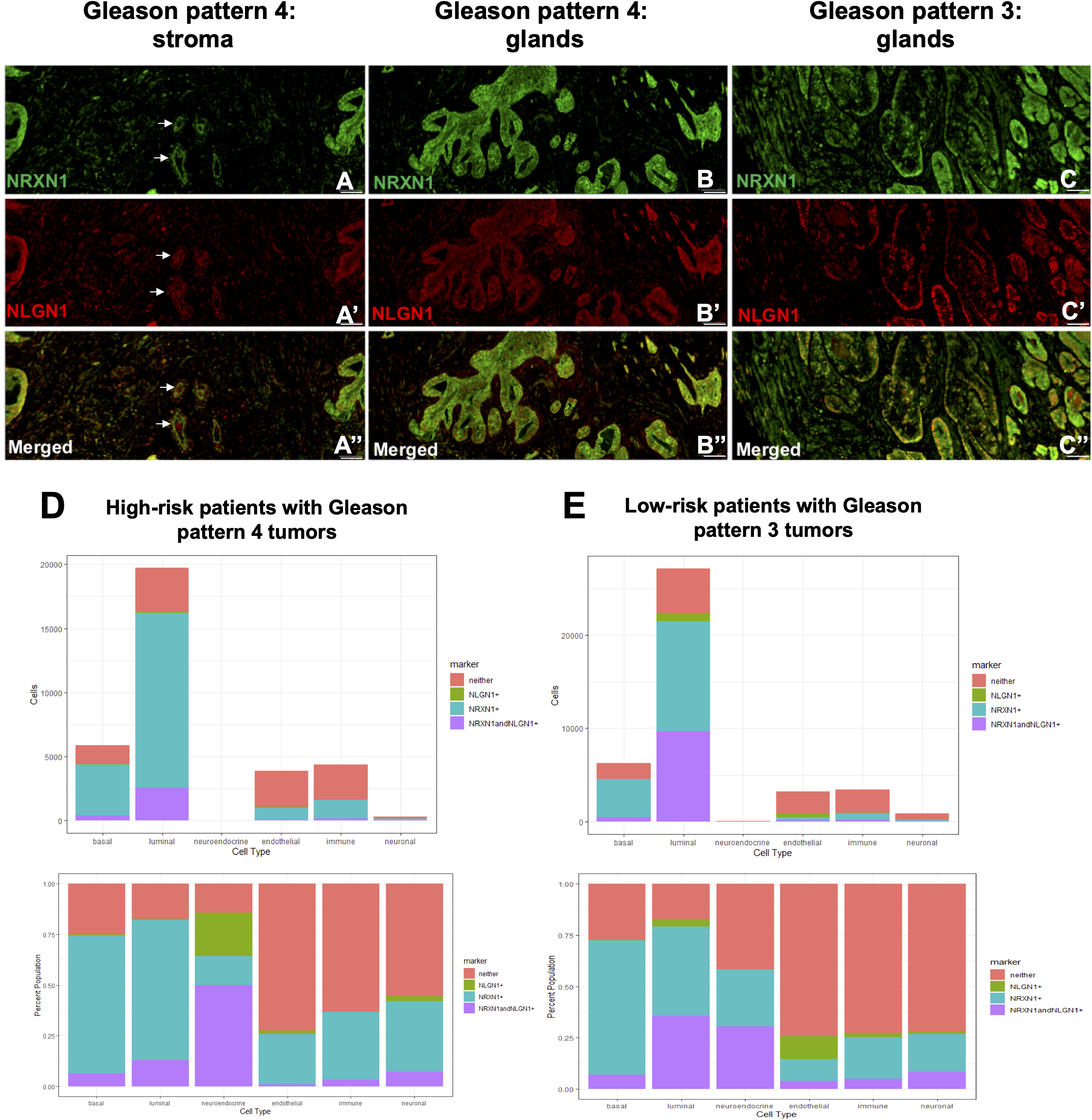
Common trans-regulatory networks exist in Gleason pattern 4 versus 3 prostate tumors. A. Transcription factor binding motifs enriched in aggregated single-cells from Gleason pattern 4 (top) and Gleason pattern 3 (bottom) tumors. Binding motifs are shown based on sequence similarities. Log scales vary due to differences in peak number and coverage. B. Transcription factor binding motifs were enriched in aggregated single-cells from three different Gleason pattern 4 tumors. C. Transcription factor binding motifs were enriched in prostate tumors from the TCGA cohort. Bulk ATAC-seq profiles from five Gleason score 3+4, six Gleason score 4+4 and ten Gleason score 4+5 patients were compared with each other.

Next, to determine if different Gleason pattern 4 tumors from different patients share similar trans-regulatory networks, we analyzed the TF binding motifs of each Gleason pattern 4 tumor individually. We observed that each Gleason pattern 4 tumor was enriched for FOXA1, HOXB13 and CDX2 binding motifs, suggesting that higher-grade prostate tumors converge on the same trans-regulatory landscape (Figure 4B). Specifically, we observed Sample_3 and Sample_4 to have a very similar TF binding motif enrichment profile, in terms of strong enrichment for FOXA1, HOXB13 and CDX2. Interestingly, Sample_15, a Gleason pattern 4 tumor, showed high enrichment for FOXA1, HOXB13 and CDX2, followed by Fra1/2, Atf3, JunB and AP-1 binding sites enriched in Gleason pattern 3 tumors, suggesting a transitionary state between the two regulatory states (Figure 4B).

We observed increased accessibility for the promoter region of TFs HOXB13 and AR in prostate tumors with Gleason pattern 4 as compared to pattern 3 (Supplementary Figure 7). FOXA1 seemed to be accessible across most prostate cancer cells and CDX2 showed higher accessibility specifically in one of the Gleason pattern 4 tumors (Supplementary Figure 7A). We also checked the patient outcome data using the Cistrome cancer database for TF profiles of FOXA1, HOXB13 and CDX2 in the TCGA PRAD data set [52, 53]. We observed that patients with PRAD tumors have higher expression of FOXA1 and HOXB13 compared to the normal prostate tissue and poor survival is associated with high FOXA1 expression (Supplementary Figure 8).

To validate the findings from our cohort, we compared the trans-regulatory landscapes of prostate tumors with different Gleason grades using the TCGA PRAD data set. In parallel to our observations, we found FOXA1, HOXB13 and CDX2 to be highly enriched in prostate tumors with Gleason pattern 4 (Gleason score 4+4 and 4+5 patients) as compared to tumors that are predominantly Gleason pattern 3 (Gleason score 3+4) (Figure 4C). Similarly, we also observed Fra1/2, Atf3, JunB and AP-1 binding sites were enriched in predominantly Gleason pattern 3 tumors as compared to higher grade prostate tumors (Figure 4C). When we analyzed the trans-regulatory differences between prostate tumors with a secondary Gleason pattern 5 (Gleason score 4+5) to tumors with Gleason pattern 4 (Gleason score 4+4) we found an enrichment of binding sites for class I steroid receptors, androgen (ARE), glucocorticoid (GRE) and progesterone (PG) (Figure 4C). Interestingly, tumors with Gleason pattern 4 still showed an enrichment for FOXA1, HOXB13 and CDX2, as well as for AP-1 like TFs as compared to tumors with a secondary Gleason pattern 5 (Figure 4C).

### Synaptic molecules NRXN1 and NLGN1 are expressed in the epithelial and stromal cells in prostate cancer

Neuronal adhesion molecules identified through our sci-ATAC-seq experiments, NRXN1 and NLGN1, are primarily expressed in the central nervous system and function in the modulation of synaptic transmission [54]. NRXN1 belongs to the family of neurexins, which is localized to the presynaptic membrane, and interacts with neuroligins such as NLGN1, which is localized to the postsynaptic membrane [55]. The expression and function of NRXN1 and NLGN1 in prostate cancer have not been characterized before. Cyclic immunofluorescent (cyclic IF) microscopy provides in-depth information about molecular composition and spatial distribution of cellular heterogeneity by allowing capture of more than 30 markers from single tissue sections [56, 57]. To determine the expression pattern of NRXN1 and NLGN1 in prostate cancer across different cell types, we performed cyclic IF staining on tissue sections from eight patients in our cohort, using cell-type specific markers. To profile the spatial heterogeneity of NRXN1 and NLGN1 expression across distinct cell types, we marked basal (CK5, CK14), luminal (CK8) and neuroendocrine (Chromogranin A) epithelial cells within the prostate glands, as well as the endothelial (CD31), neuronal (NCAM) and immune cells (CD45, CD3) in the prostate cancer microenvironment (Figure 5, Supplementary Figure 9). We validated antibodies against NRXN1 and NLGN1 using a brain tissue section as positive and colon tissue as negative control [58, 59] (Supplementary Figure 9).

**Figure 5:**
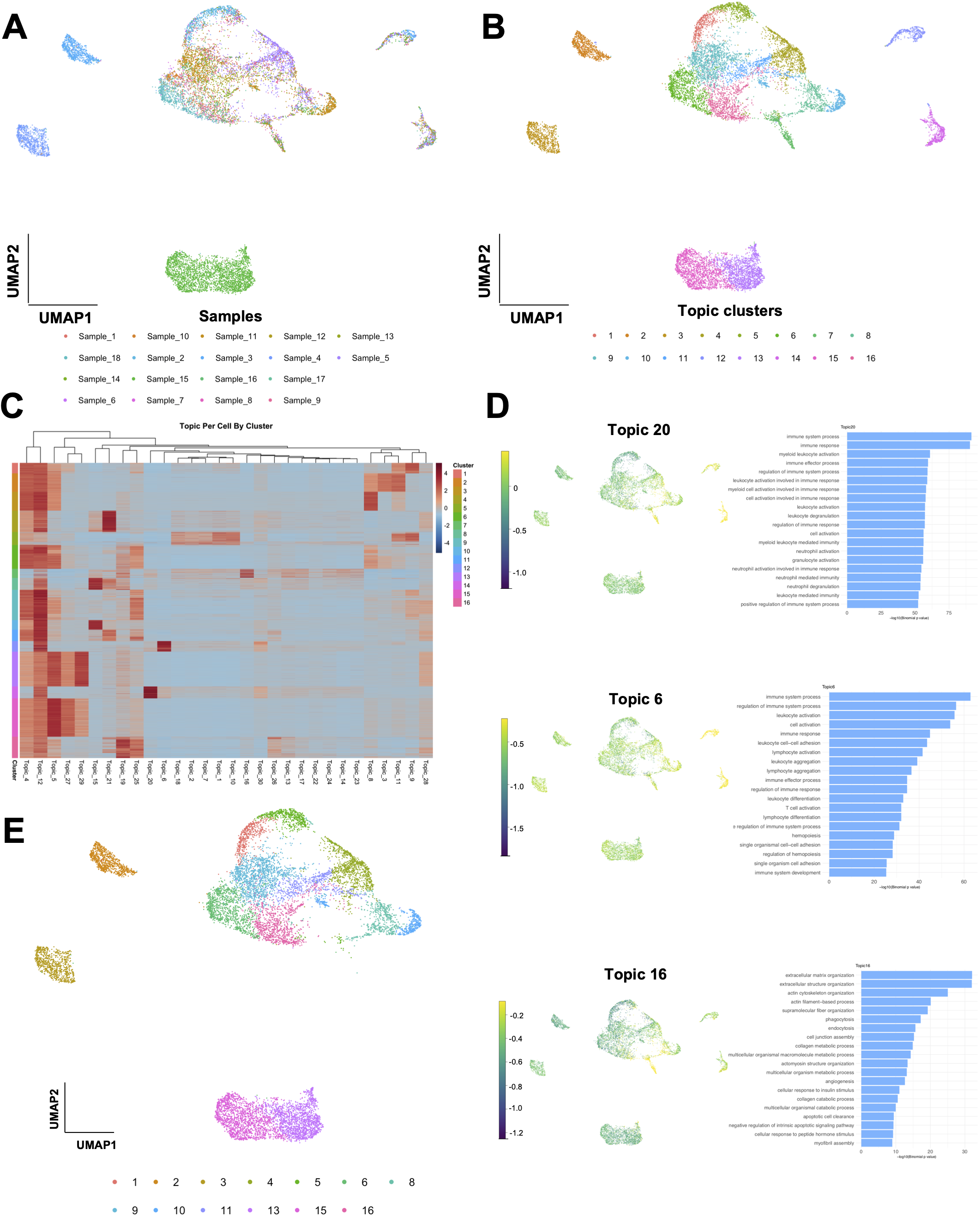
Neuronal adhesion molecules NRXN1 and NLGN1 are expressed in the epithelial and stromal cells in prostate cancer. A-C. Protein expression of NRXN1 (green) and NLGN1 (red) in tumors with Gleason pattern 4 (Sample_6) (A-B) and 3 (Sample_8) (C) and tumor stroma (Sample_6). Supplementary Figure 10 shows H&E images and ROIs for each sample. White arrows point to expression of NRXN1 and NLGN1 in the blood vessels (A). Scale bars (white line) show 200 pixels. D. Expression of NRXN1 and NLGN1 across different cell types from three tissue sections acquired from high-risk patients with Gleason pattern 4 prostate tumors in our cohort. X-axis shows cell types and Y-axis shows the total number of segmented cells (top) or percentage population of cells positive for that marker (bottom). NRXN1 expressing cells are shown in blue and NLGN1 in green. Cells that express both proteins are shown in purple and cells that do not express either are marked with pink. E. Expression of NRXN1 and NLGN1 across different cell types from five tissue sections acquired from low-risk patients with Gleason pattern 3 tumors in our cohort. X-axis shows cell types and Y-axis shows the total number of segmented cells (top) or percentage population of cells positive for that marker (bottom). NRXN1 expressing cells are shown in blue and NLGN1 in green. Cells that express both proteins are shown in purple and cells that do not express either are shown in pink.

We found that both NRXN1 and NLGN1 were expressed in basal, luminal and neuroendocrine cells in the prostate glands (Figure 5D-E, Supplementary Figure 10). We observed that there was a slight increase in the number of neuroendocrine cells that expressed NRXN1 and NLGN1 in tissue sections from high-risk patients (Figure 5D-E). We observed NRXN1 and NLGN1 were expressed in blood vessels in the prostate cancer tissue sections (Figure 5A-D-E). Both our sci-ATAC-seq data and our cyclic IF results indicated NRXN1 and NLGN1 expression in immune cells infiltrating the prostate tumors (Figure 3E, Figure 5D-E). Finally, we observed that both NRXN1 and NLGN1 were expressed in the N-CAM positive neuronal cells within the prostate tumor stroma, with overall slightly higher expression profiles in high-risk patients (Figure 5D-E).

## DISCUSSION

Prostate cancer exhibits significant heterogeneity at the molecular, cellular and tissue level, hampering efforts to accurately determine risk of progression for localized prostate cancer. Single-cell technologies provide unique opportunities to accurately delineate the stages of prostate tumor heterogeneity prior to metastasis. However, single-cell technologies can also generate large data sets that may be difficult to validate and mechanistically resolve at the context of the tissue. Therefore, it is crucial to merge these single-cell sequencing technologies with spatial imaging to provide an understanding of the transitionary states between indolent and aggressive cancers.

Using sci-ATAC-seq, we identified a co-accessibility pattern in neuronal adhesion molecules, NRXN1 and NLGN1, that distinguishes Gleason pattern 4 prostate tumors from Gleason pattern 3. Recent evidence shows that both the peripheral nervous system and progenitors from the central nervous system may influence prostate cancer progression [60–62]. Neuronal cells within the prostate cancer microenvironment can release neurotransmitters that may modulate the behavior of prostate cancer cells [63, 64]. It has been shown that non-neuronal cells expressing neurexins and neuroligins result in pre- and post-synaptic specialization in neurons and these calcium-dependent synaptic molecules can exploit calcium channels present in surrounding cells for their biological activity [65–67]. More interestingly, NLGN1 has shown to be cleaved enzymatically at its N-terminal ectodomain and secreted at excitatory synapses by enzymatic cleavage [68, 69]. Similarly, NLGN3 has been shown to be secreted in an activity-dependent manner in glioma, promoting cell division and glioma growth [70]. Our cyclic IF analysis revealed the spatial distribution of NRXN1 and NLGN1 in prostate cancer and identified cell-type specific expression patterns for both proteins. We observed prostate epithelial cells, as well as immune and neuronal cells express these synaptic molecules in prostate cancer. Based on our findings, we propose that expression of neuronal adhesion molecules in prostate cancer cells mark tumors for a more aggressive, potentially metastatic phenotype. Our findings suggest the possibility that NRXN1 and NLGN1 expression and secretion in high-grade prostate tumors may modulate the activity between cancer cells and neurons, which may contribute to perineural invasion or neoneurogenesis in prostate cancer [62, 71, 72].

Our study has provided an atlas of the regulatory landscape of low-grade (Gleason pattern 3) and high-grade (Gleason pattern 4) prostate tumors. Interestingly, we did not observe regulatory heterogeneity among single-cells from Gleason pattern 4 tumors. This may be because aggressive prostate tumors acquire distinct evolutionary trajectories that involve different types of chromosomal arrangements [73–75]. For instance, we observed at least one of the Gleason pattern 4 tumors had high levels of accessibility for the ERG promoter that led to a very distinct chromatin accessibility landscape for cells from this tumor. On the other hand, Gleason pattern 3 tumors exhibited significant cell-to-cell heterogeneity that led to clusters of cells from multiple Gleason pattern 3 tumors. Interestingly, all cells from Gleason pattern 3 tumors were bound by chromatin restraints that were lost in Gleason pattern 4 tumors. Further genomic studies are necessary to provide links between the common chromosomal rearrangements and single-cell epigenomic states of prostate cancer cells.

Our results also indicate unique trans-regulatory signatures for different grade localized prostate tumors. We found low-grade prostate tumors were significantly enriched for the AP-1 family of TF binding sites (JUN, JUNB, JUND and FOS, FOSB and FRA1, and the closely related activating TFs: ATF and CREB), in contrast to highgrade prostate tumors that were enriched for FOXA1, HOXB13 and CDX2. Interestingly, we observed at least one of the high-grade prostate tumors possessed signatures for both transcriptional regulatory programs. We do not know whether this tumor represents a case that is undergoing a transition in chromatin structure, i.e. the low-grade prostate tumor regulatory marks are still present even though the tumor transitioned to a more aggressive state.

We also observed that all Gleason pattern 4 tumors in our cohort share the same trans-regulatory circuit, even though they formed distinct clusters of cells based on their chromatin accessibility profiles. Strikingly, we were able to identify the same trans-regulatory signature in Gleason pattern 4 and higher tumors in the TCGA data set. Our results, combined with the analysis of the TCGA data set, show that prostate tumors with Gleason pattern 4 share a common transcriptional regulatory program defined by an enrichment of FOXA1, HOXB13 and CDX2 binding sites. This finding is particularly interesting since prostate cancers usually have low numbers of recurrent mutations [48, 74, 76]. Future research will determine whether disparate genomic changes observed in prostate cancer converge on common epigenomic profiles and if these present druggable targets in non-metastatic tumors.

Identification of gene regulatory programs shared by high- and low-grade tumors present a strong opportunity to identify biomarkers for patient stratification despite the overwhelming molecular and cellular heterogeneity that exists in prostate cancer. There are several drugs in development targeting FOXA1 and HOXB13 [77–79]. Our results suggest that these therapies could potentially benefit patients with high-grade non-metastatic prostate tumors. However, further mechanistic studies are required to better evaluate the potential effect of these drugs on chromatin structure and transcriptional regulation in localized tumors.

It is unclear whether the enriched binding sites for FOXA1, HOXB13 and CDX2 are dependent or independent of AR activity in single cells from high-grade tumors. It is known that FOXA1, HOXB13 and GATA2 act as pioneer TFs in the prostate tissue to facilitate androgen receptor (AR) transcription during prostate carcinogenesis [80]. However, their AR-independent activity has not been studied extensively. We anticipate that there is a similar synergy between FOXA1, HOXB13 and CDX2 TFs in remodeling the chromatin structure in high-grade prostate tumors. Previous studies show colocalization of TFs FOXA1 and HOXB13 at the reprogrammed AR binding sites in human prostate cancer cells, and draw a link between TMPRSS2-ERG fusion and cooption of FOXA1 and HOXB13 to specific regulatory elements across the genome [81]. Interestingly, we observed an enrichment for FOXA1 and HOXB13 in Gleason pattern 4 tumors independent of their ERG accessibility (Supplementary Figure 10). However, it is not possible to infer any details about the cooperation of these TFs with each other, their synergy with AR, and their functional impact on transcription based on our results. Additional characterization of histone modification markers is necessary to understand the details of the synergy between these TFs on downstream gene expression in Gleason pattern 4 vs 3 prostate tumors.

## METHODS

### Isolation of prostate adenocarcinoma cells from radical prostatectomy samples

Radical prostatectomy samples from 18 patients were obtained from the Biolibrary at OHSU. Fresh-frozen prostate samples positive for prostate adenocarcinoma were recovered from each radical prostatectomy specimen. H&E images were reviewed by a pathologist to confirm the Gleason grade and score of each sample, as well as score assigned by the Biolibrary. Five unstained adjacent 5-micron tissue sections were obtained from each sample for immunofluorescent microscopy studies.

### Risk stratification of the patient cohort

The Cancer of the Prostate Risk Assessment Post-Surgical (CAPRA-S) score and post-radical prostatectomy nomogram from Memorial Sloan Kettering Cancer Center (MSKCC) were used to identify patients with low-, intermediate- and high-risk prostate cancer [32, 35]. CAPRA-S score is calculated based on factors such as pathological Gleason score, surgical margin status and presence or absence of extracapsular extension and lymph node involvement for patients who have gone through radical prostatectomy surgery to estimate the probability of disease recurrence [33].

We also analyzed bulk ATAC-seq profiles of 21 patients available through the TCGA data set. We calculated the CAPRA scores for these patients and found that the majority of them were either intermediate- or high-risk as opposed to our cohort of patients that consisted mainly of low- or high-risk patients based on CAPRA risk stratification (Supplementary File 2).

### Whole-mount sample acquisition

Radical prostatectomy specimens were obtained via standard of care surgery and processed in the OHSU Pathology lab. Subjects consented to study OHSU IRB #18321. Under this protocol an annotated radiology worksheet depicting the location of the MRI-detected tumor foci was completed by the radiologist and a copy was sent with the specimen to the pathologist and grossing technician. Fresh prostate specimens were inked and sectioned at 5 mm intervals using a prostate slicing device (Procut P/5, Milestone Medical), followed by a gross examination to correlate gross findings with the annotated radiology worksheet. Fresh tissue was collected from tumor area with a 5 mm punch biopsy and removed for research purposes. A frozen section from this research tissue sample was used to confirm the presence of tumor. The entire remaining prostate was then submitted and processed for whole mount histology. Formalin fixed paraffin embedded (FFPE) tissue blocks and slides for standard clinical annotation were generated. This work was done under the supervision of a subspecialized genitourinary pathologist.

### sci-ATAC-seq Sample Processing

#### Sample preparation and nuclei isolation

The nuclei isolation protocol was improved and barcode space was extended to increase multiplexing ability of the combinatorial indexing protocol. Frozen prostate tissue samples (0.1-0.8 grams) were homogenized in Nuclei Isolation Buffer (NIB, 10 mM TrisHCl pH7.4, 10 mM NaCl, 3 mM MgCl2, 0.1% Igepal, 1 protease inhibitor tablet (Roche, Cat. 11873580001)) using a dounce homogenizer. Isolated nuclei were washed three times with ice cold 1XPBS and centrifuged down at 500 x g for 5 min at 4^°^C. Washed nuclei were passed through a 35μm cell strainer (Corning) and stained with 5 μL (5 mg/ml) DAPI to mark the nuclei.

#### Tn5 transposome assembly

Tn5 enzyme was purified as described previously and loaded with specific oligo sequences [82]. Tn5 adaptor sequences synthesized at Integrated DNA technologies (Supplementary File 3). Briefly, oligonucleotides for ME-rev (phosphorylated 19-basepair mosaic end) and i5 or i7 were incubated in equimolar amounts (100 μM each) for 5 min at 95^°^C and cooled down slowly on the thermocycler in 3^°^C increments. 0.25vol adaptor sequences, 0.4 vol 100% glycerol, 0.12 vol 2X dialysis buffer (100 mM HEPES–KOH at pH 7.2, 0.2 M NaCl, 0.2 mM EDTA, 2 mM DTT, 0.2% Triton X-100, 20% glycerol), 0.1 vol Tn5 (50 μM), 0.13 vol water. Uniquely indexed transposomes were stored in 96-well plates at −20^°^C.

#### Cell sorting

For the first sort plate, 3000 DAPI stained nuclei were sorted into 96-well plates using the FacsAria cell sorter. Each well contained 10 μL tagmentation buffer (5 μL NIB and 5 μL TD buffer from Illumina). For the second sort plate, 22-25 DAPI stained nuclei were sorted into 96-well plates using FacsAria cell sorter. Each well contained 8.5 μL master mix (0.25 μL 20mg/ml BSA, 0.5 μL 1% SDS, 7.5 μL distilled water and 2.5 μL i5 and i7 PCR index primers).

#### Tagmentation

Tagmentation reaction was carried out at 55°C for 30 minutes after the 1^st^ nuclei sort. Once the reaction was at room temperature the plates were placed on ice and samples from each well were pooled in an Eppendorf tube. Pooled nuclei were passed through a 35μm cell strainer (Corning) and stained with 5 μL (5 mg/ml) DAPI before the second nuclei sort. Samples were incubated at 55^°^C for 15 min to denature the transposase after the second sort.

#### PCR indexing

Nuclei were amplified using RT-PCR for 20-25 cycles to insert unique i5-i7 DNA oligo sequences in each well based on the previously published protocol [50] (Supplementary File 3). 7.5 μL NPM PCR mix (Illumina), 4 μL distilled water, 0.5 μL 100X SYBR Green dye was added to each well and the following PCR amplification cycle was followed: 75^°^C for 5 min, 98^°^C for 30 sec, (for 22-25 cycles) 98^°^C for 10 sec, 63^°^C for 30 sec, 72^°^C for 60 sec, plate read at 72^°^C for 10 sec.

#### Sample purification and fragment analysis

5 μL of sample from each well was pooled from each well and the library pool was purified using Qiagen PCR purification kit followed by AMPure bead purification. Each library pool was analyzed using Bioanalyzer to assess the quantity and distribution of fragment size before sequencing. Libraries were sequenced on the Next-seq platform (Illumina) using a 150-cycle kit with a custom sequencing recipe (Read 1: 47 imaged cycles; Index 1: 8 imaged cycles, 27 non-imaged / dark cycles, 10 imaged cycles; Index 2: 8 imaged cycles, 21 non-imaged / dark cycles, 10 imaged cycles; Read 2: 47 imaged cycles) [7].

### Immunohistochemistry

Each formalin fixed and paraffin embedded (FFPE) prostate tissue block was serially sectioned. One Hematoxylin-and-Eosin (H&E) stained section and five adjacent unstained sections with 5-micron thickness were acquired from all tumors. FFPE tissue sections were deparaffinized as previously described [56]. Each tissue section was incubated in a 65^°^C oven for one hour and then immediately transferred in Xylene solution. Slides were sequentially immersed in fresh Xylene solution two times, 5 minutes each; 100% ethanol two times 5 minutes each; 95% ethanol two times 2 minutes each; 70% ethanol two times 2 minutes each and distilled water two times 5 minutes each. Antigen retrieval was done in a medical decloaking chamber filled with 0.5L distilled water. Tissue slides were placed in a plastic jar that contains Citrate buffer (pH=6) and incubated at high pressure for 15 minutes. Each slide was then dipped in distilled water and incubated in pH9 buffer for 15 minutes before being transferred to distilled water at room temperature to complete antigen retrieval. Slides were blocked with 10% NGS, 1%BSA in PBS for 30 minutes at room temperature inside a humidity chamber. Primary antibodies conjugated to Alexa Fluor dyes were prepared in 5% NGS, 1%BSA in PBS buffer. Conjugated primary antibodies used in this study are shown in Supplementary Figure 9 and in the key resources table. All slides were imaged using Zeiss Axioscan microscope located at the Knight Cancer Institute Research Building.

### Cyclic Immunofluorescence Microscopy

For each round of immunofluorescent staining primary antibodies conjugated to Alexa Fluor dyes 488, 555, 647 and 750 were mixed in 1% BSA, 5% NGS solution. Slides were incubated either for 2 hours at room temperature or overnight at 4^°^C and washed four times in 1X PBS for 5 minutes. Tissue sections were mounted in Slowfade Gold DAPI mounting media and imaged using a Zeiss Axioscan microspcope. After a successful scan was obtained, the fluorophore signal was quenched via incubating slides in a quenching solution (10% 10X PBS, 0.4% 5M NaOH, 3% H_2_O_2_) under broad spectrum light for an hour. Slides were washed three times in 1X PBS for 5 minutes and mounted in Slowfade Gold DAPI mounting media and imaged with Axioscan to confirm quenching of the signal. We acquired images of each slide to measure the autofluorescence levels at rounds 3 and 6 (Supplementary Figure 9).

### Antibody Conjugation

Buffer exchange has been done using Amicon ultra 10 KDa spin columns for antibodies that had sodium azide as preservative. Alexa Fluor dyes were prepared by dissolving each dye in DMSO to a final concentration of 10 mM. Each antibody was mixed with 1M NaHCO_3_ in 10:1 volume ratio. 0.6 μl of Alexa Fluor dye was added per 100 μg of antibody. Conjugation reaction was carried out on a rocker for 2 hours at room temperature in dark. A buffer exchange using Amicon ultra 10 KDa spin columns was performed to remove the excess Alexa Fluor dye.

### Cell lines

PC3 (ATCC CRL-1435) and LNCaP (ATCC FGC CRL-1740) human prostate epithelial cells were maintained in RPMI 1640 medium with 10% FBS at 37°C and 5% CO2 at recommended densities. Adherent cells were detached using TrypLE Express (Gibco) and were collected at mid-log phase for all experiments. After collection, cells were washed twice with ice cold 1X PBS. Cells were then filtered with a 35-μm cell strainer (Corning). Cell viability and concentration were measured with Trypan blue on the Countess II FL (Life Technologies). Cell viability was greater than 90% for all samples.

### Cell line fixation, staining and imaging

PC3 cells and LnCAP cells were plated at 20k/well and 40k/well concentrations respectively on 96 well plates with #1.5 polymer coverslip bottoms (Cellvis). After reaching 80% confluence, cells were washed once with 1X DPBS(-) and fixed with 4% PFA diluted from a freshly opened 16% PFA (Electron Microscopy Sciences) ampoule, for 15 minutes at RT. Fixed cells were washed with 1X DPBS(-) once and permeabilized and blocked in a 3% BSA solution (Thermo Fisher) with 0.5% Triton-X-100 (Sigma-Aldrich). Primary antibodies were prepared at 1:50 – 1:100 dilution in 3% BSA solution and incubated with cells for 2 hrs at RT or overnight at 4^°^C. If needed, secondary antibodies were diluted in 3% BSA and incubated for 1hr at RT, followed by application of 0.01mg/mL DAPI in 1X PBS (Thermo Fisher) (or 5mg/ml diluted 1:500) for 10 minutes at RT. After each staining step cells were washed with 1X DPBS(-) three times. Cells were imaged using a Zeiss/Yokogawa CSU-X1 spinning disk confocal setup with a 40X objective.

### sci-ATAC-seq Data Analysis and Visualization

#### Read Alignment and Pre-processing

Analysis of reads were done using snapATAC (Single Nucleus Analysis Pipeline for ATAC-seq) [40]. Bases were converted to fastq format using bcl2fastq. Reads were then aligned using snaptools aligned-paired-end command using all of the default parameters. Reads were aligned to hg19 using the – bwa parameter which utilizes the bwa aligner [83]. Reads were than preprocessed using the snaptools snap-pre command using the default parameters: --min-mapq=30 --min-flen=0 --max-flen=1000 --keep-chrm=TRUE --keep-single=TRUE --keep-secondary=False --max-num=1000000 --min-cov=100.

#### sci-ATAC-seq Binned Counts Matrix and Peak Counts Matrix Generation and quality control (QC)

Two count matrices were used in this study, one being a binned matrix where each counts were generated for each 5kb bin in the genome. This matrix was used for clustering. The binned counts were generated using the snaptools -- snap-add-bmat command. For the peak matrix, the R package snapATAC command runMACS was used using the MACS2 [84] parameters: --nomodel --shift 100 --ext 200 --qval 5e-2 -B –SPMR. The peak matrix was then generated and added using the createPmat function in the snapATAC.

#### Single-Cell Clustering and Visualization

The matrix of binned counts was used binarized and inputted into a latent dirichlet allocation dimensionality reduction utilizing the method described by the tool cisTopic [41]. This was done using the snapATAC runLDA function using 30 topics. The Uniformed Manifold Approximation and Projection (UMAP) algorithm was then applied to the top 30 topics using the snapATAC runUMAP function. To further cluster the cells, the runCluster function was applied utilizing the Louvain method for community network detection [85].

#### Differentially Accessible Regions Analysis

Differential accessibility was performed on the MACS2 callpeak matrix using the findDARs function in snapATAC. This function utilizes the edgeR package using the exactTest method [86]. P values were than adjusted using the Benjamini-Hochberg method. Two differentially accessible lists were generated for each comparison, one with pvalues < 0.05 and another with FDR <0.05.

#### GREAT Analysis

To further understand the epigenetic factors in cell groups, the top 1500 peak regions per topic were saved and fed into Genomic Regions Enrichment of Annotations Tool (GREAT). Further, differentially accessible regions from individual comparisons were also used as input to GREAT [47].

#### HOMER Analysis

Motif enrichment was done using the homer findMotifsGenome.pl command using a bed file of interest with the parameters: -size 200 -mask [51].

#### Cicero Analysis

Cis-regulatory interactions and co-accessibility scores were plotted with the R package using the function plot_connections [49].

### Image registration and background subtraction

Raw TIFF image files from each round of cyclic IF imaging were registered using the nuclear DAPI image from the first round as reference. Background subtraction was done using a blank imaging cycle with no fluorescently tagged antibodies to remove the tissue autofluorescence signal from the images using a matrix subtraction operation [87].

### Single-cell segmentation, feature extraction and quantification

DAPI image from the very last round of imaging was used to account for the tissue loss over cycles. Nuclei and cytoplasm segmentation were done using the QiTissue software with the following parameters: Nuclei segmentation method (Advanced Morphology for Tissue), Use nuclei cycle (8), Detection Sensitivity (100%), Min/Max Diameter (28/29 pixels), Separation Force (100%), Cytoplasm segmentation method (Donut), Detection Sensitivity (100%), Max Diameter (150 pixels), Perinuclear region (3 pixels), Donut width (10 pixels), Neighbor Touch Region (8 pixels).

Average signal intensity of each marker in nucleic and cytoplasmic compartments, nucleic x and y coordinates, nucleus and cell size features were extracted for each segmented cell using QiTissue’s “Measure Cell Features” options.

Cells with abnormal size features that did not fall within approximately the 5-to-95 percentile range were assumed to be incorrectly segmented, either multiple cells merged together or a fragment of the cell segmented, and excluded from the analysis.

Three regions of interest (ROIs) were selected from each tissue section based on the H&E images reviewed by a pathologist. ROIs were selected to avoid areas of tissues with edge effects or staining gradients. Within each ROI thresholds for NRXN1, NLGN1 and different cell type markers were determined. Using the thresholds for cell type markers, we identified luminal (CK8), basal (CK5, CK14) and neuroendocrine (Chromogranin A) epithelial cells, immune cells (CD3), endothelial cells (CD31) and neuronal cells (N-CAM). Based on the cell type and marker (NRXN1 and NLGN1) thresholds, we identified cells that express only NRXN1, only NLGN1, both NRXN1 and NLGN1 or neither for each cell type. The quantities of each of these categories for each cell type was plotted using the “ggplot2” library of R, version 3.4.3.

### Survival Analysis

We used the Cancer Transcription Factor Targets pipeline within the Cistrome Cancer (“the set of cis-acting targets of a trans-acting factor on a genome-wide scale, also known as the in vivo genome-wide location of [transcription factor binding-sites] or histone modifications”) [52]. Survival Analysis was plotted using the TCGA PRAD data set, selecting for transcription factors FOXA1, HOXB13 and CDX2.

## ACKNOWLEDGEMENTS

We are grateful to all patients who participated to the OHSU study (IRB #18321) that provided tissue from prostatectomy whole-mounts. R.P.K was funded by the Collins Medical Trust for this study. We are also grateful to the Biolibrary and the Histopathology Shared Resource (HSR) at OHSU for providing patient samples, specifically Aletha Letsch and Cheyenne Martin. The Biolibrary and HSR was supported by NIH grants P30 CA069533 and P30 CA069533 13S5 through the Knight Cancer Institute. We thank Erin Watson for collecting patient information. We thank the Spellman lab and the prostate cancer working group in CEDAR for engaging discussions. We are grateful for the collaborative environment in the Knight Cancer Institute and productive interactions with the Gray, the Chang and the Adey labs. We thank Kemal Sonmez for his input on single-cell data analysis. We thank Crystal Shaw from the Advanced Light Microscopy Core and Dorian LaTocha from the Flow Cytometry Core at OHSU for their technical help. We thank Shelley Barton, Hisham Mohammed, Joshua Saldivar, Stefanie Linch and Lindsey Minter for providing feedback on the manuscript. S.E.E was funded by a full CEDAR Grant (“Investigating the epigenetic heterogeneity and tumor microenvironment in prostate tumors at single-cell resolution) for this study.

## AUTHOR CONTRIBUTIONS

S.E.E., A.A. and P.T.S. conceived the study. S.E.E. performed all experiments with help from Z.S. and A.F. sci-ATAC-seq data analysis was performed by A.C. and A.A., while A.C. and S.E.E. performed the analysis of TCGA data. Z.S. and S.E.E. performed analysis of cyclic IF data. G.T. and R.P.K. recruited patients and collected whole-mount patient samples. G.V.T. reviewed the pathology. S.E.E. processed all patient samples. S.E.E., Z.S., A.C., G.T., A.A., P.T.S performed the biological analysis and interpretation. S.E.E. wrote the manuscript with input from all authors.

## MATERIALS & CORRESPONDANCE

Further information and requests for resources and reagents should be directed to Sebnem Ece Eksi.

### Materials Availability

All unique/stable reagents generated in this study are available when accompanied by a completed Materials Transfer Agreement.

### Data and Code Availability

The accession numbers for the sci-ATAC-seq reported in this paper are GEO: (submitted, to be determined). The raw single-cell ATAC-sequencing files and processed data files generated in this study are available in GEO under the super series GSE (submitted, to be determined). Single-cell combinatorial indexing data is available under GSE (submitted, to be determined).

Code used for the analysis of sci-ATAC-seq data in this study is available on Github (https://github.com/AlexChitsazan/ProstateTumorATACCode). Code used for cyclic IF analysis in this study is available on GitHub (https://github.com/zeynepsayar/Neuronal_cyclic_R). Any other scripts of analysis can be requested from the authors.

## SUPPLEMENTAL FIGURE TITLES AND LEGENDS

**Supplementary Figure 1: 14,251 cells were sequenced from 18 prostate tumor samples.** A. Number of cells sequenced per sample is shown. B. Distribution of cells from each tumor sample into different clusters. C. All DNA libraries were sequenced to near saturation as shown by median duplication rates. Quality measurements of the sci-ATAC-seq libraries produced in the study are shown, TotalNumberOfBarcodes: Total Number of Cells, MedianNumOfSequencingFragments: Median Number of Sequencing Fragments Per Cell, MedianNumberOfUniquelyMappedFragments: MedianNumberOfUniquelyMappedFragments, MedianNumberOfMappabilityRatio:

Median Mapping Ratio Per Cell, MedianNumberOfProperlyPairedRatio: Median Ratio of Reads With A Paired Read Per Cell, MedianNumberOfDuplicateRatio: Median Ratio of Duplicates Per Cell, MedianNumberOfChrMRatio: Median Ratio of ChrM Mapped Reads Per Cell, Sample: Sample cistopic Name, SampleNamePaper: Sample Name Paper. D. 16 Clusters that were identified around 30 Topics exhibit hierarchical clustering as shown by a cluster dendogram. Clusters 12 and 14 consisted of immune cells, which formed a separate cluster in the dendogram. Cluster 7 consisted of stromal cells associated with the prostate cancer epithelial cells.

**Supplementary Figure 2: The top differentially accessible chromatin regions identified in prostate tumors with Gleason pattern 3 versus 4.** A. Top 23 accessible regions in Gleason pattern 4 tumors are marked with red in the plot. B. The top 279 differentially accessible chromatin regions are identified in prostate tumors with Gleason pattern 3 versus 4 (marked with red). C. Cells that have chromatin accessibility around the promoter of four genes, PDE7B, WNT5A, SLC44A5 and HMGN2P46, selected from the list of top 23 regions are shown in red on the UMAP. D. GREAT GO results for accessible DNA elements in prostate tumors with Gleason pattern 3.

**Supplementary Figure 3: Distinct Topics exist in Clusters that define tumors with Gleason pattern 3 and 4.** Topics 27 and 29 are found to be enriched in Clusters 13 and 15, and Topics 8, 3 and 11 are enriched in Clusters 2, 3 and 4, which together define the accessible chromatin regions of prostate tumors with Gleason pattern 4 (Gleason score 4+3, 4+4 and 4+5). Topics 15 and 21 are enriched in prostate tumors with Gleason pattern 3 (Gleason score 3+3 and 3+4). Topics 4, 12, 19 and 25 are found to be associated with all prostate tumors. A. Heatmap showing the distribution of topics across different Gleason scores identified common and distinct pathways associated with Gleason pattern 3 and 4 tumors. B. GREAT GO results for Topics 4, 12, 29, 27, 21 and 15 that are associated with Clusters defining prostate tumor subgroups in the study.

**Supplementary Figure 4: Gleason pattern 3 cells do not form patient-specific clusters when analyzed independently from Gleason pattern 4 cells.** A. Single-cells from prostate samples were clustered using cisTOPIC based on their chromatin accessibility profiles. Cells are colored according to their sample IDs on the UMAP. B. Single-cells from prostate samples were clustered using cisTOPIC based on their chromatin accessibility profiles. Cells are colored according to their cluster IDs on the UMAP.

**Supplementary Figure 5: Chromatin accessibility profiles of genes associated with known prostate cancer subtypes are enriched in specific clusters.** A. ERG, ETV1, ETV4 and FLI1 accessibility are shown in red. Samples 4 and 18 show high accessibility for ERG whereas Sample 3 showed high accessibility to FLI1 and ETV1. B. PTEN, SPOP, MYC and IDH1 accessibility are shown in red. Higher accessibility for MYC is observed in tumors with Gleason pattern 4 (outer class). C-D. Cicero maps show higher number of putative regulatory interactions around the CDH9 (C) and NLGN1 (D) loci in Gleason pattern 3 (top) vs 4 (bottom) tumors.

**Supplementary Figure 6: NRXN1 and CDH9 are significantly accessible in tumors with Gleason pattern 4 in the TCGA bulk ATAC-seq data set.** A. Distribution of Gleason scores across patients are shown in the TCGA bulk ATAC-seq data set. x-axis shows the number of patients in the cohort, y-axis indicates the Gleason score. B. Distribution of Gleason scores across patients are shown in the TCGA RNA-seq data set. C. GREAT GO results (top) for TCGA bulk ATAC-seq data for prostate tumors with Gleason pattern 4 (Gleason score 4+4) as compared to tumors predominantly Gleason pattern 3 (Gleason score 3+4). Identified peaks that are linked to CDH9 and NRXN1 are shown (bottom). D. GREAT GO results (top) for TCGA bulk ATAC-seq data for prostate tumors with predominantly Gleason pattern 4 (Gleason score 4+5) as compared to tumors predominantly Gleason pattern 3 (Gleason score 3+4). Identified peaks that are linked to CDH9 and NRXN1 are shown (bottom). E. NRXN1 is expressed at the RNA level in PRAD tumors from TCGA. Bulk RNA-seq results from 497 patients are shown, each dot representing a patient. Patients are categorized into columns based on their Gleason scores as indicated on the x-axis. Fragments Per Kilobase of transcript per Million mapped reads (FPKM) are shown on the y-axis. A. NRXN1 expression is shown across patients with different Gleason scores. F. NLGN1 expression is shown across patients with different Gleason scores. G. GAPDH (control) expression is shown across patients with different Gleason scores.

**Supplementary Figure 7: Chromatin accessibility profiles of specific transcription factors are enriched in prostate tumors with Gleason pattern 3 and 4. Cells that have accessibility for the promoter region of the gene of interest are shown in red.** A. Accessibility to the promoter region of TFs FOXA1, HOXB13, CDX2 and AR are shown. B. Accessibility to the promoter region of JUN, FOS, ATF3 and FOSL1 are shown.

**Supplementary Figure 8: Patient outcome data from the TCGA data set shows the link between the expression levels of TFs FOXA1, HOXB13 and CDX2 (left), which are expressed in the tumor (black bar) and normal (red bar) tissue.** Patient survival (right) data is shown for the top 25^th^ percentile of patients (red line) and the bottom 25^th^ percentile (black line).

**Supplementary Figure 9: Antibodies against NRXN1 and NLGN1 were tested and validated using different tissue sections and cell lines for cyclic IF.** A. Brain tissue section stained with the NLGN1 (green) and NRXN1 (red) as a positive control for the antibodies. White arrows show an example region with positive NLGN1 and NRXN1 expression. Scale bar (white line) shows 200 pixels. B. Colon tissue section stained with the NLGN1 (green) and NRXN1 (red) as a negative control for the antibodies. Scale bar (white line) shows 200 pixels. C. NRXN1 antibody tested on fixed PC3 prostate cancer cells (C, magenta) shown in contrast to DAPI (C’, blue). C’’ shows the two channels merged. White arrow shows NRXN1 expression. Scale bar (white line) shows 20 μm. D. NRXN1 antibody tested on fixed LNCaP prostate cancer cells (D, magenta) shown in contrast to DAPI (D’, blue). D’’ shows the two channels merged. White arrow shows NRXN1 expression. Scale bar (white line) shows 20 μm. E. NLGN1 antibody tested on fixed LNCaP prostate cancer cells (E, magenta) shown in contrast to DAPI (E’, blue). E’’ shows the two channels merged. White arrow shows NLGN1 expression. Scale bar (white line) shows 20 μm. F. Rounds table for cyclic IF experiments show cell-type specific markers. Each column is marked by the Alexa Fluor (AF) dye that are attached to the antibodies within that column: AF488, AF555, AF647 and AF750. Each row shows the antibodies that are used in that round of antibody staining. Autofluorescence (AF) levels are measured at the end of round 2 and round 5.

**Supplementary Figure 10: NRXN1 and NLGN1 are expressed in tumors with Gleason pattern 3 and 4.**

H&E tissue sections from all patients are shown (left). Three regions of interest (ROIs) were drawn in each whole-mount tissue section based on their H&E staining (blue diagrams). The expression of NRXN1 and NLGN1 are measured across different cell types within each ROI (bar graphs). x-axis shows cell types and y-axis shows the total number of segmented cells or percentage population of cells positive for that marker. NRXN1 expressing cells are shown in purple and NLGN1 in green. Cells that express both proteins are shown in blue and cells that do not express either are marked with pink.

**Supplementary File 1: CAPRA-S and MSKCC nomogram calculations of patients in our cohort.**

**Supplementary File 2: CAPRA calculations of patients in the TCGA PRAD cohort.**

**Supplementary File 3: Tn5 and PCR oligo sequences used in the study.**

