## Supplementary Figures 1-10 for "Single-cell analysis of localized low- and high-grade prostate cancers"

A

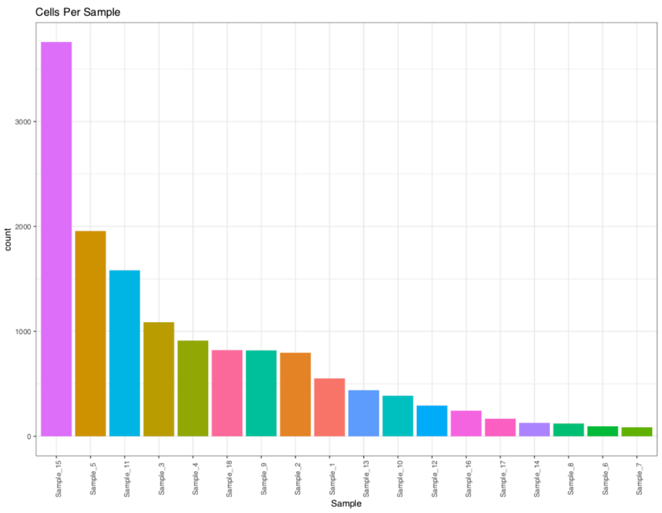


B

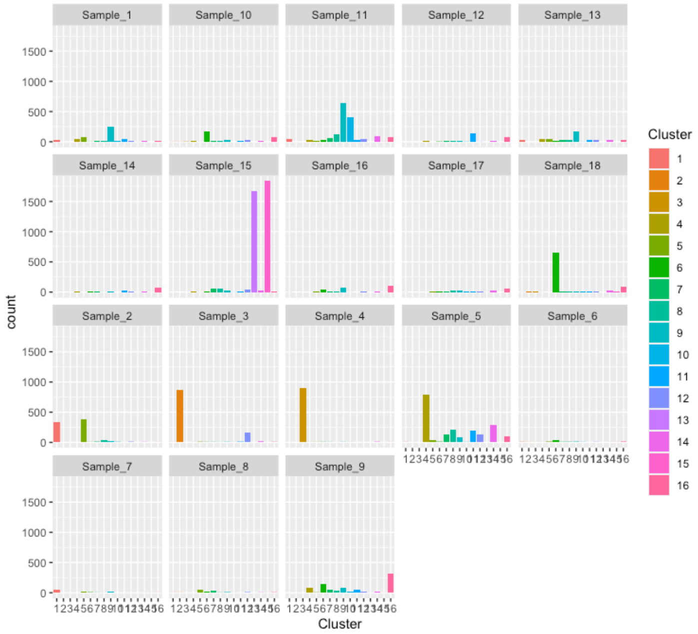


C

**Descriptive statistics**

| *SampleNamePaper* | *Row* | *TotalNumberOfBarcodes* | *MedianNumOfSequencingFragments* | *MedianNumberOfUniquelyMappedFragments* | *MedianNumberOfMappabilityRatio* | *MedianNumberOfProperlyPairedRatio* | *MedianNumberOfDuplicateRatio* | *MedianNumberOfChrMRatio* |
|---|---|---|---|---|---|---|---|---|
| NA | AllSamples December.Ida | 14242 | 13436 | 1287.5 | 0.87 | 1 | 0.86 | 0.02 |
| Sample_1 | G33_1 | 551 | 30893 | 1102 | 0.86 | 1 | 0.96 | 0.05 |
| Sample_2 | G33_2 | 796 | 10402.5 | 1998 | 0.89 | 1 | 0.78 | 0.03 |
| Sample_3 | G4_1 | 1087 | 70051 | 1803 | 0.91 | 1 | 0.97 | 0.02 |
| Sample_4 | G4_2 | 912 | 22114 | 3374 | 0.89 | 1 | 0.84 | 0.01 |
| Sample_7 | ID54773 | 86 | 3740.5 | 725 | 0.87 | 1 | 0.76 | 0.1 |
| Sample_6 | ID56893 | 95 | 8705 | 648 | 0.88 | 1 | 0.92 | 0.09 |
| Sample_5 | G33_3 | 1956 | 13263.5 | 3337 | 0.88 | 1 | 0.72 | 0.04 |
| Sample_8 | TB12146 | 122 | 34111.5 | 640.5 | 0.78 | 0.88 | 0.97 | 0.02 |
| Sample_9 | TB4732 | 819 | 7759 | 898 | 0.82 | 1 | 0.86 | 0.03 |
| Sample_10 | TB5020 | 387 | 9242 | 1209 | 0.87 | 1 | 0.86 | 0.02 |
| Sample_11 | TB6475 | 1582 | 7102.5 | 825 | 0.87 | 1 | 0.87 | 0.02 |
| Sample_12 | TB7521 | 293 | 5766 | 665 | 0.87 | 1 | 0.86 | 0.04 |
| Sample_13 | TB7677 | 439 | 18007 | 2229 | 0.83 | 1 | 0.87 | 0.1 |
| Sample_14 | TB7799 | 128 | 39672.5 | 714 | 0.8 | 0.87 | 0.98 | 0.02 |
| Sample_15 | TB9679 | 3757 | 10760 | 1358 | 0.88 | 1 | 0.87 | 0.01 |
| Sample_16 | WM306009 | 244 | 42762 | 673.5 | 0.77 | 0.88 | 0.98 | 0.05 |
| Sample_17 | WM306010 | 167 | 33514 | 644 | 0.79 | 0.89 | 0.97 | 0.03 |
| Sample_18 | WM306016 | 821 | 38961 | 704 | 0.81 | 0.89 | 0.97 | 0.04 |

D

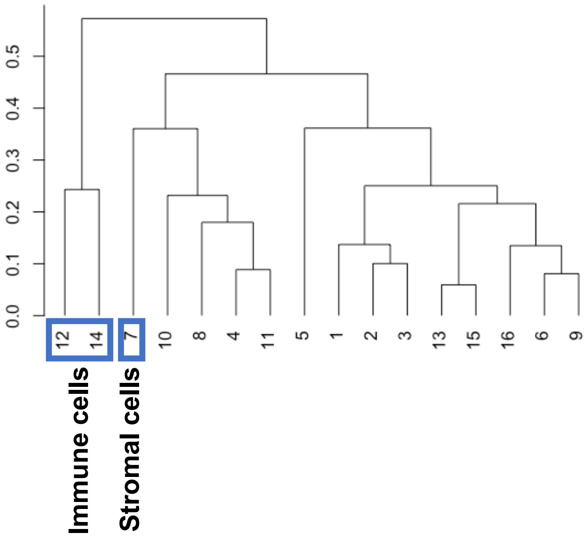

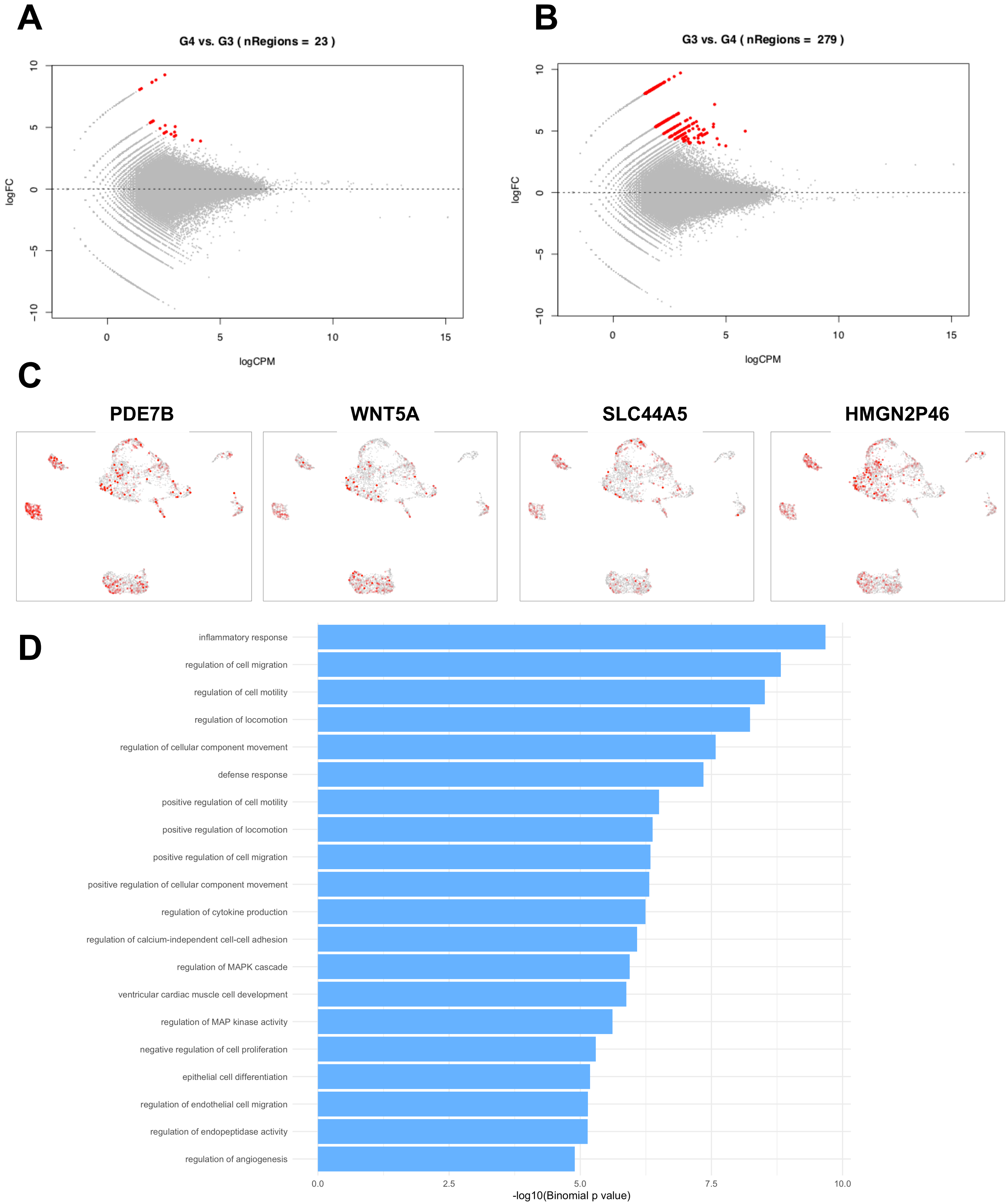

A
G4 vs. G3 ( nRegions = 23 )
logFC
logCPM
B
G3 vs. G4 ( nRegions = 279 )
logFC
logCPM
C
PDE7B
WNT5A
SLC44A5
HMGN2P46
D
inflammatory response
regulation of cell migration
regulation of cell motility
regulation of locomotion
regulation of cellular component movement
defense response
positive regulation of cell motility
positive regulation of locomotion
positive regulation of cell migration
positive regulation of cellular component movement
regulation of cytokine production
regulation of calcium-independent cell-cell adhesion
regulation of MAPK cascade
ventricular cardiac muscle cell development
regulation of MAP kinase activity
negative regulation of cell proliferation
epithelial cell differentiation
regulation of endothelial cell migration
regulation of endopeptidase activity
regulation of angiogenesis
-log10(Binomial p value)

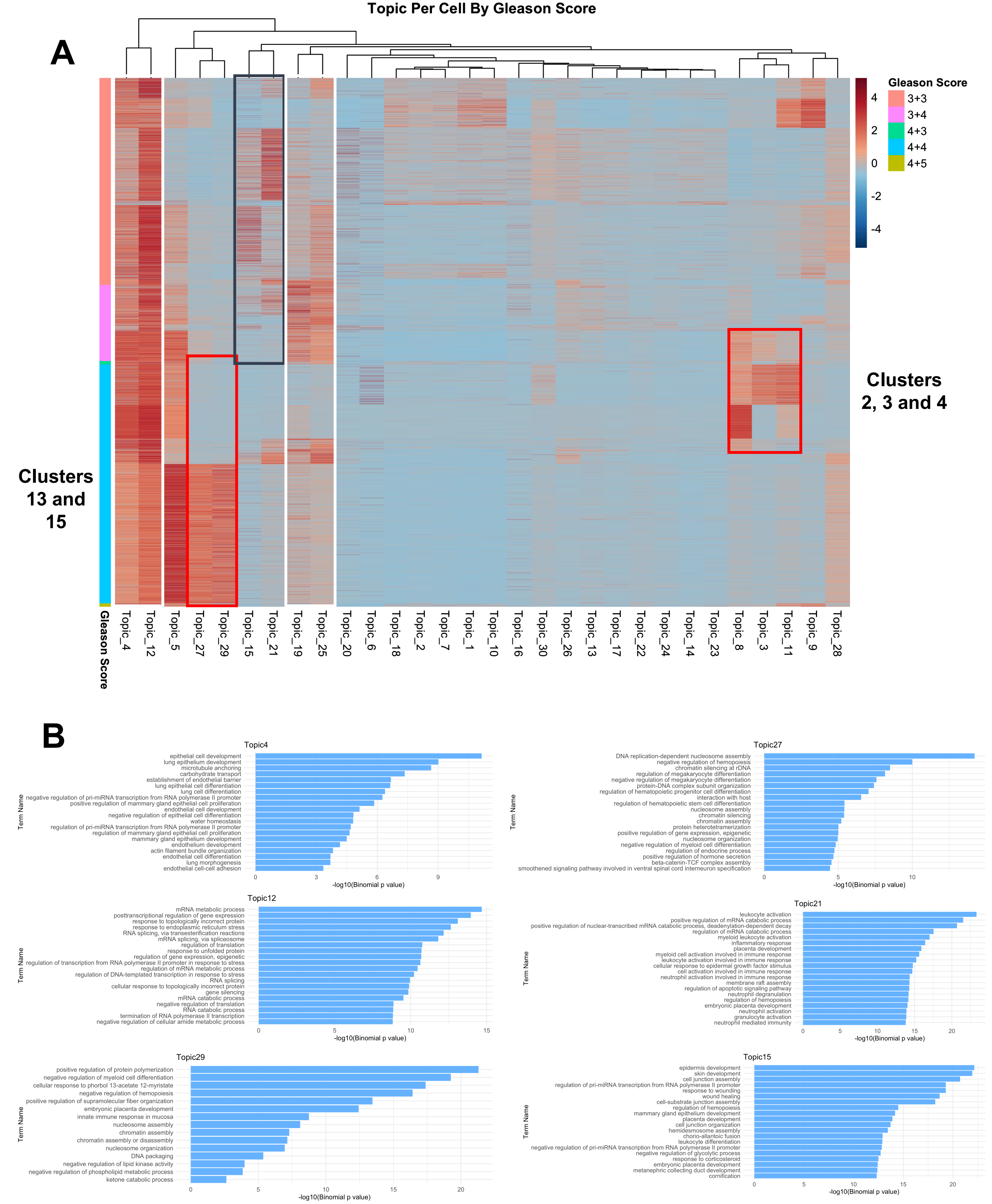

Topic Per Cell By Gleason Score
A
Gleason Score
3+3
3+4
4+3
4+4
4+5
4
2
0
-2
-4
Clusters 2, 3 and 4
Clusters 13 and 15
Gleason Score
Topic_4
Topic_12
Topic_5
Topic_27
Topic_29
Topic_15
Topic_21
Topic_19
Topic_25
Topic_20
Topic_6
Topic_18
Topic_2
Topic_7
Topic_1
Topic_10
Topic_16
Topic_30
Topic_26
Topic_13
Topic_17
Topic_22
Topic_24
Topic_14
Topic_23
Topic_8
Topic_3
Topic_11
Topic_9
Topic_28
B
Topic4
epithelial cell development
lung epithelium development
microtubule anchoring
carbohydrate transport
establishment of endothelial barrier
lung epithelial cell differentiation
lung cell differentiation
negative regulation of pri-miRNA transcription from RNA polymerase II promoter
positive regulation of mammary gland epithelial cell proliferation
endothelial cell development
negative regulation of epithelial cell differentiation
water homeostasis
regulation of pri-miRNA transcription from RNA polymerase II promoter
regulation of mammary gland epithelial cell proliferation
mammary gland epithelium development
endothelium development
actin filament bundle organization
endothelial cell differentiation
lung morphogenesis
endothelial cell-cell adhesion
Term Name
-log10(Binomial p value)
Topic27
DNA replication-dependent nucleosome assembly
negative regulation of hemopoiesis
chromatin silencing at rDNA
regulation of megakaryocyte differentiation
negative regulation of megakaryocyte differentiation
protein-DNA complex subunit organization
regulation of hematopoietic progenitor cell differentiation
interaction with host
regulation of hematopoietic stem cell differentiation
nucleosome assembly
chromatin silencing
chromatin assembly
protein heterotetramerization
positive regulation of gene expression, epigenetic
nucleosome organization
negative regulation of myeloid cell differentiation
regulation of endocrine process
positive regulation of hormone secretion
beta-catenin-TCF complex assembly
smoothened signaling pathway involved in ventral spinal cord interneuron specification
Term Name
-log10(Binomial p value)
Topic12
mRNA metabolic process
posttranscriptional regulation of gene expression
response to topologically incorrect protein
response to endoplasmic reticulum stress
RNA splicing, via transesterification reactions
mRNA splicing, via spliceosome
regulation of translation
response to unfolded protein
regulation of gene expression, epigenetic
regulation of transcription from RNA polymerase II promoter in response to stress
regulation of mRNA metabolic process
regulation of DNA-templated transcription in response to stress
RNA splicing
cellular response to topologically incorrect protein
gene silencing
mRNA catabolic process
negative regulation of translation
RNA catabolic process
termination of RNA polymerase II transcription
negative regulation of cellular amide metabolic process
Term Name
-log10(Binomial p value)
Topic21
leukocyte activation
positive regulation of mRNA catabolic process
positive regulation of nuclear-transcribed mRNA catabolic process, deadenylation-dependent decay
regulation of mRNA catabolic process
myeloid leukocyte activation
inflammatory response
placenta development
myeloid cell activation involved in immune response
leukocyte activation involved in immune response
cellular response to epidermal growth factor stimulus
cell activation involved in immune response
neutrophil activation involved in immune response
membrane raft assembly
regulation of apoptotic signaling pathway
neutrophil degranulation
regulation of hemopoiesis
embryonic placenta development
neutrophil activation
granulocyte activation
neutrophil mediated immunity
Term Name
-log10(Binomial p value)
Topic29
positive regulation of protein polymerization
negative regulation of myeloid cell differentiation
cellular response to phorbol 13-acetate 12-myristate
negative regulation of hemopoiesis
positive regulation of supramolecular fiber organization
embryonic placenta development
innate immune response in mucosa
nucleosome assembly
chromatin assembly
chromatin assembly or disassembly
nucleosome organization
DNA packaging
negative regulation of lipid kinase activity
negative regulation of phospholipid metabolic process
ketone catabolic process
Term Name
-log10(Binomial p value)
Topic15
epidermis development
skin development
cell junction assembly
regulation of pri-miRNA transcription from RNA polymerase II promoter
response to wounding
wound healing
cell-substrate junction assembly
regulation of hemopoiesis
mammary gland epithelium development
placenta development
cell junction organization
hemidesmosome assembly
chorio-allantoic fusion
leukocyte differentiation
negative regulation of pri-miRNA transcription from RNA polymerase II promoter
negative regulation of glycolytic process
response to corticosteroid
embryonic placenta development
metanephric collecting duct development
cornification
Term Name
-log10(Binomial p value)

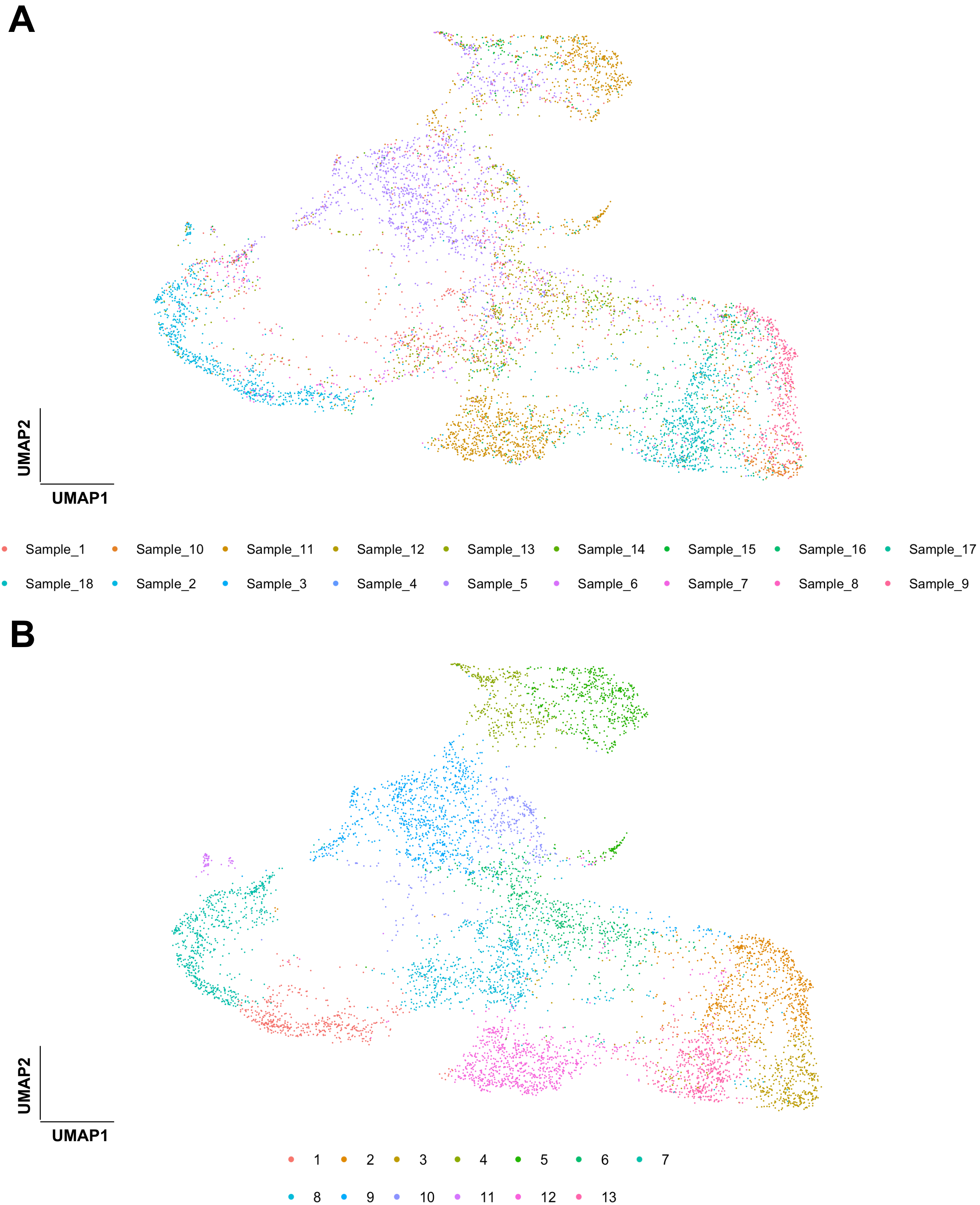
A
UMAP2
UMAP1
Sample_1
Sample_10
Sample_11
Sample_12
Sample_13
Sample_14
Sample_15
Sample_16
Sample_17
Sample_18
Sample_2
Sample_3
Sample_4
Sample_5
Sample_6
Sample_7
Sample_8
Sample_9
B
UMAP2
UMAP1
1
2
3
4
5
6
7
8
9
10
11
12
13

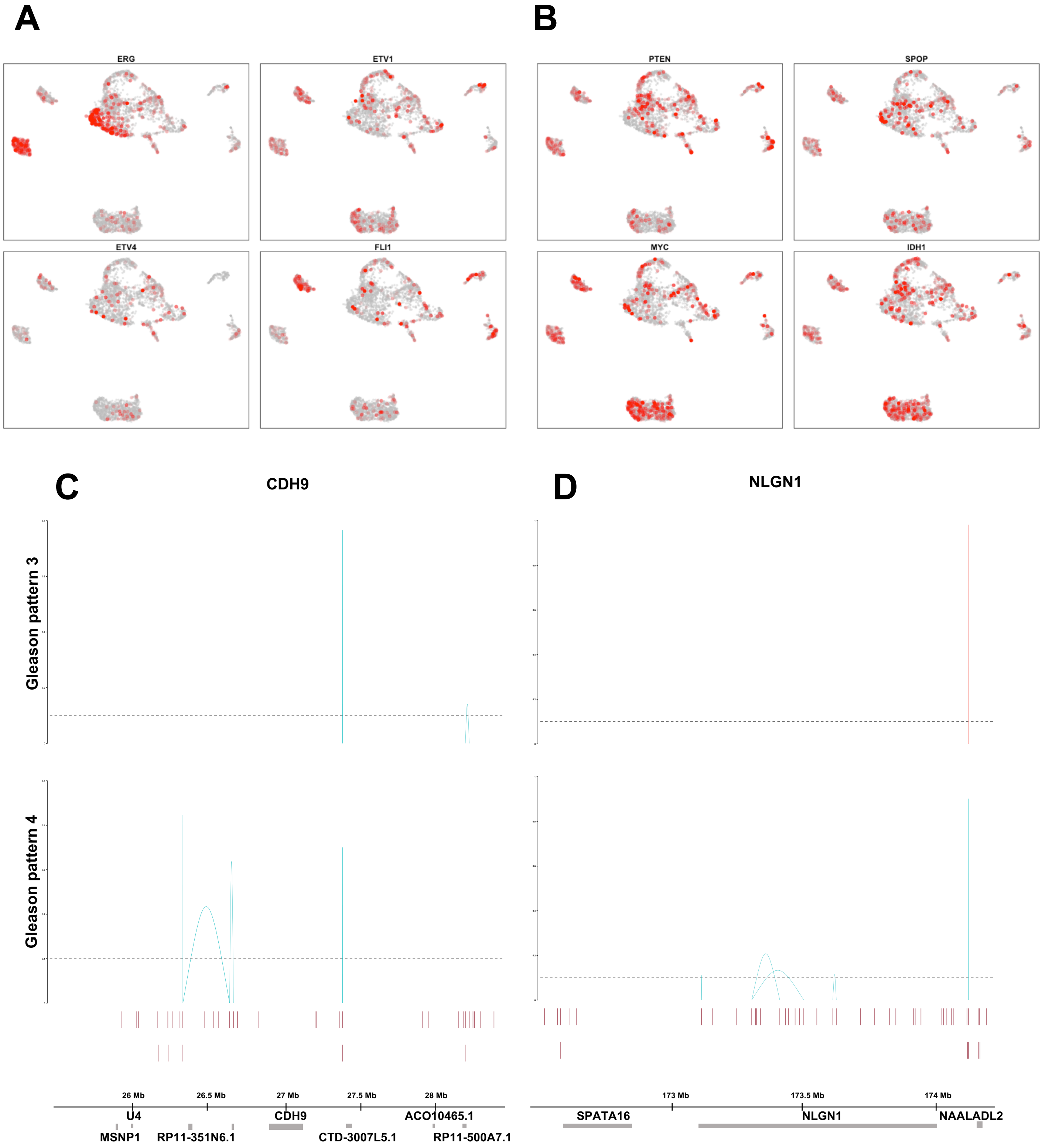
A
B
ERG
ETV1
ETV4
FLI1
PTEN
SPOP
MYC
IDH1
C
CDH9
D
NLGN1
Gleason pattern 3
Gleason pattern 4
26 Mb
26.5 Mb
27 Mb
27.5 Mb
28 Mb
U4
CDH9
AC010465.1
MSNP1
RP11-351N6.1
CTD-3007L5.1
RP11-500A7.1
173 Mb
173.5 Mb
174 Mb
SPATA16
NLGN1
NAALADL2

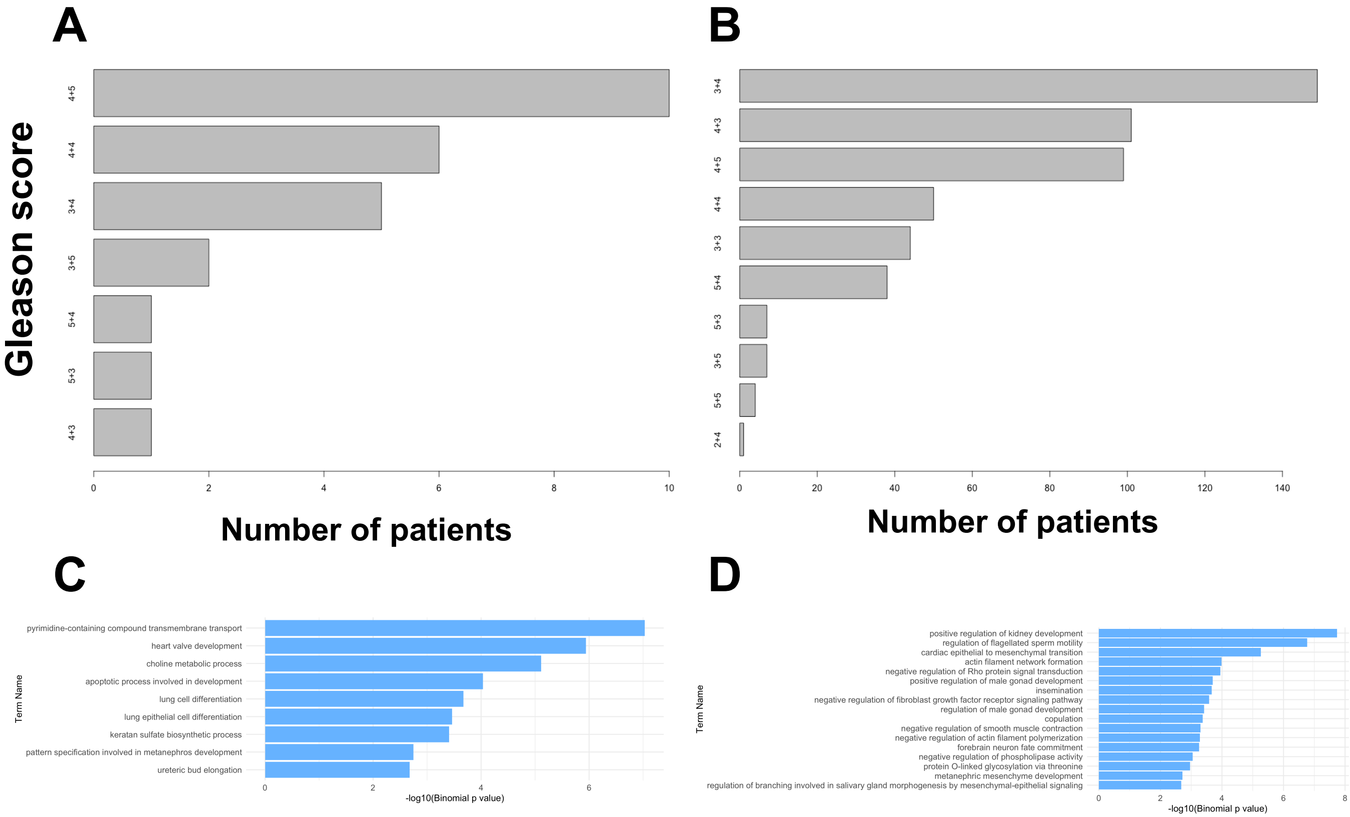


| Gene | Region (distance to TSS) |
|---|---|
| **CDH9** | -349,114 |
| **NRXN1** | -299,192, +635,988, +695,594, +989,463 |

| Gene | Region (distance to TSS) |
|---|---|
| **CDH9** | -868,089, -641,977, -349,114 |
| **NRXN1** | -732,627, -33,053, +695,594, +989,463 |

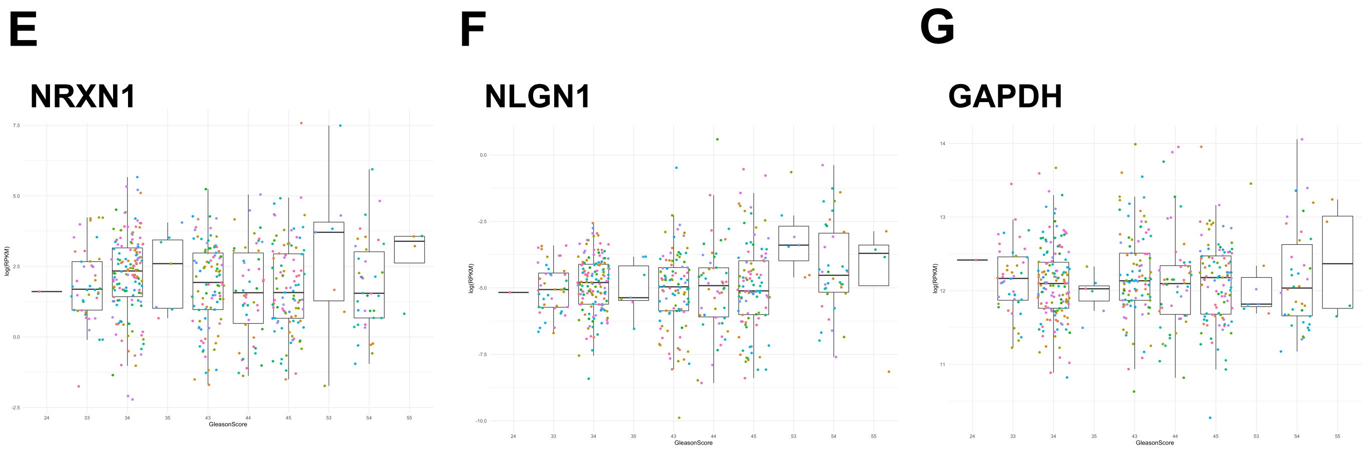

A

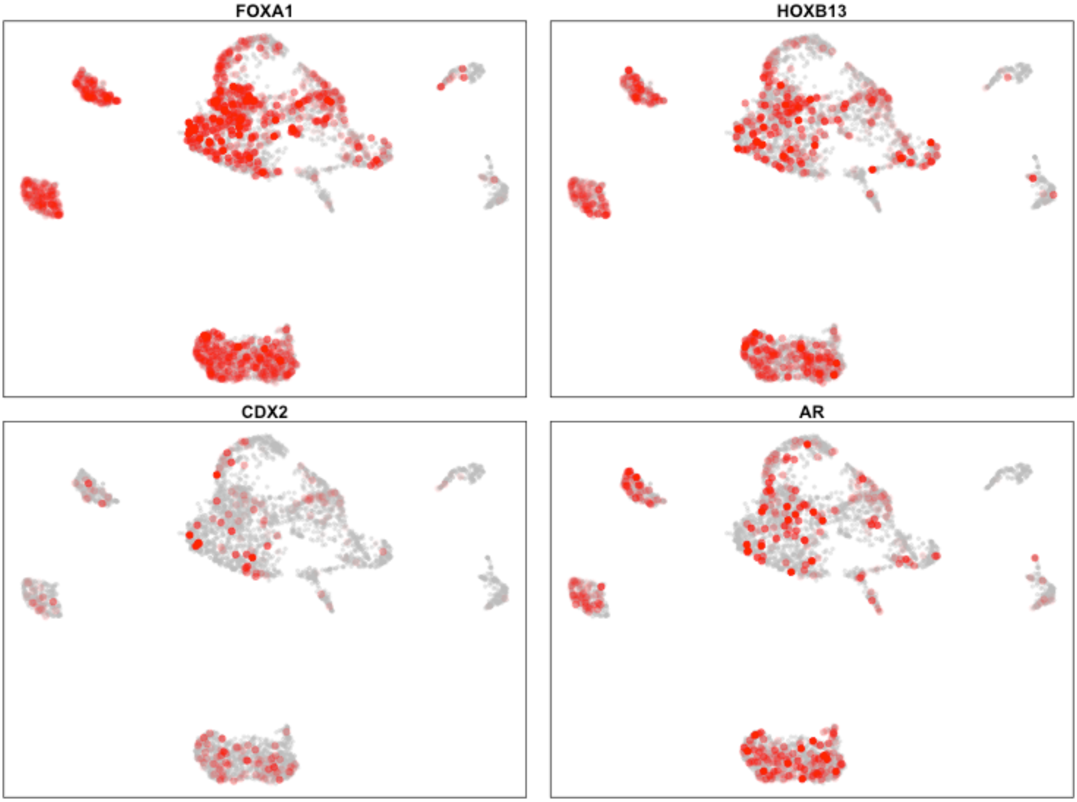


B

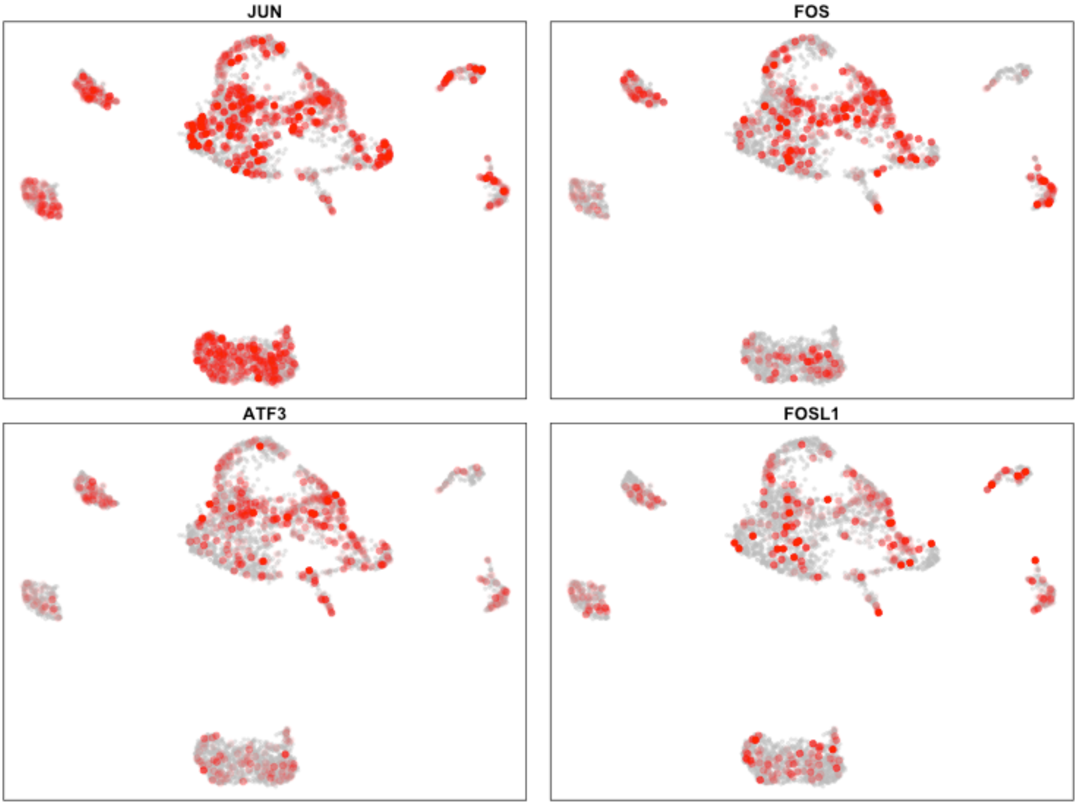

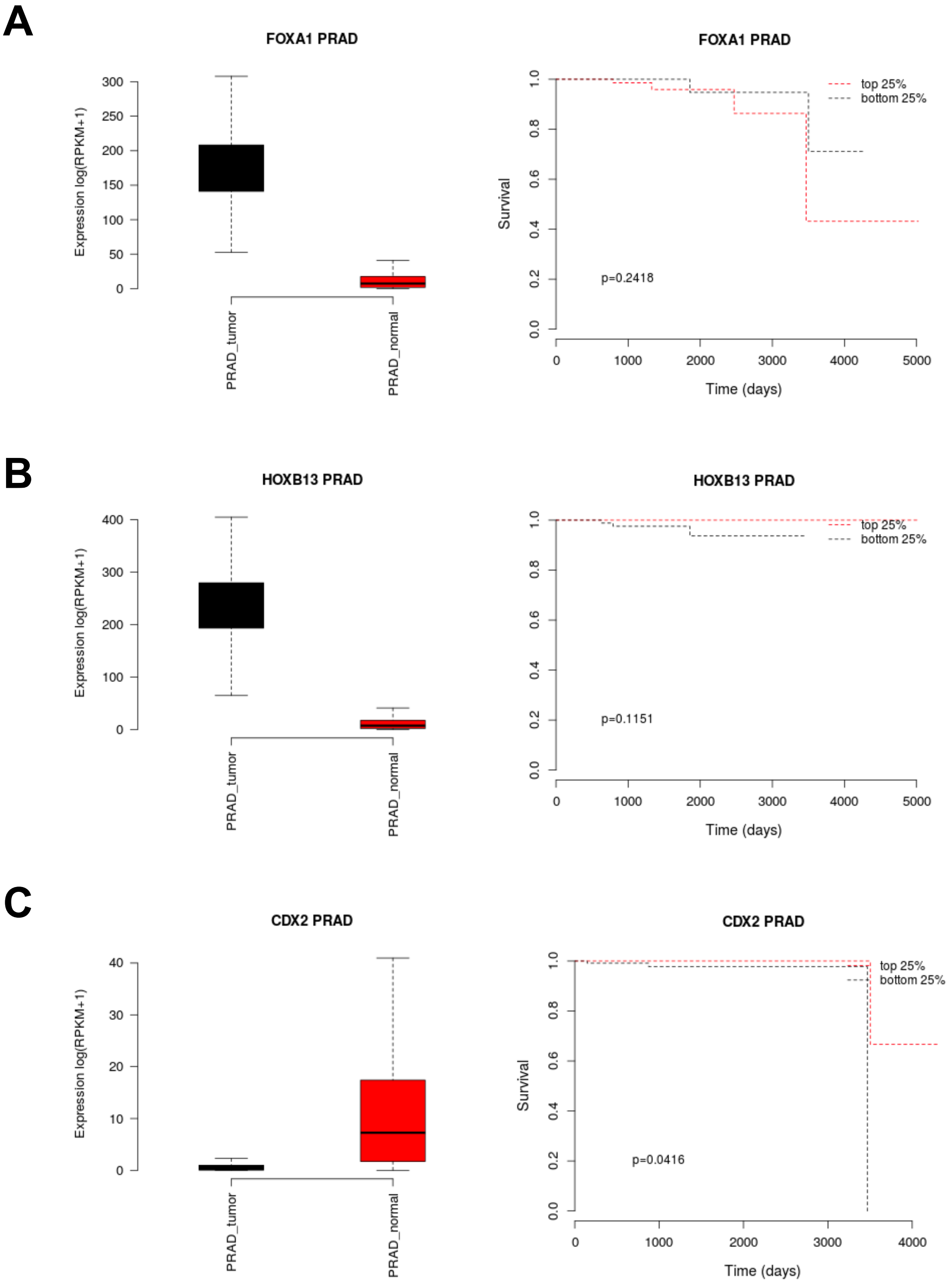

A
FOXA1 PRAD
Expression log(RPKM+1)
300
250
200
150
100
50
0
PRAD_tumor
PRAD_normal
FOXA1 PRAD
Survival
1.0
0.8
0.6
0.4
0.2
0.0
top 25%
bottom 25%
p=0.2418
0
1000
2000
3000
4000
5000
Time (days)
B
HOXB13 PRAD
Expression log(RPKM+1)
400
300
200
100
0
PRAD_tumor
PRAD_normal
HOXB13 PRAD
1.0
0.8
0.6
0.4
0.2
0.0
top 25%
bottom 25%
p=0.1151
0
1000
2000
3000
4000
5000
Time (days)
C
CDX2 PRAD
Expression log(RPKM+1)
40
30
20
10
0
PRAD_tumor
PRAD_normal
CDX2 PRAD
Survival
1.0
0.8
0.6
0.4
0.2
0.0
top 25%
bottom 25%
p=0.0416
0
1000
2000
3000
4000
Time (days)

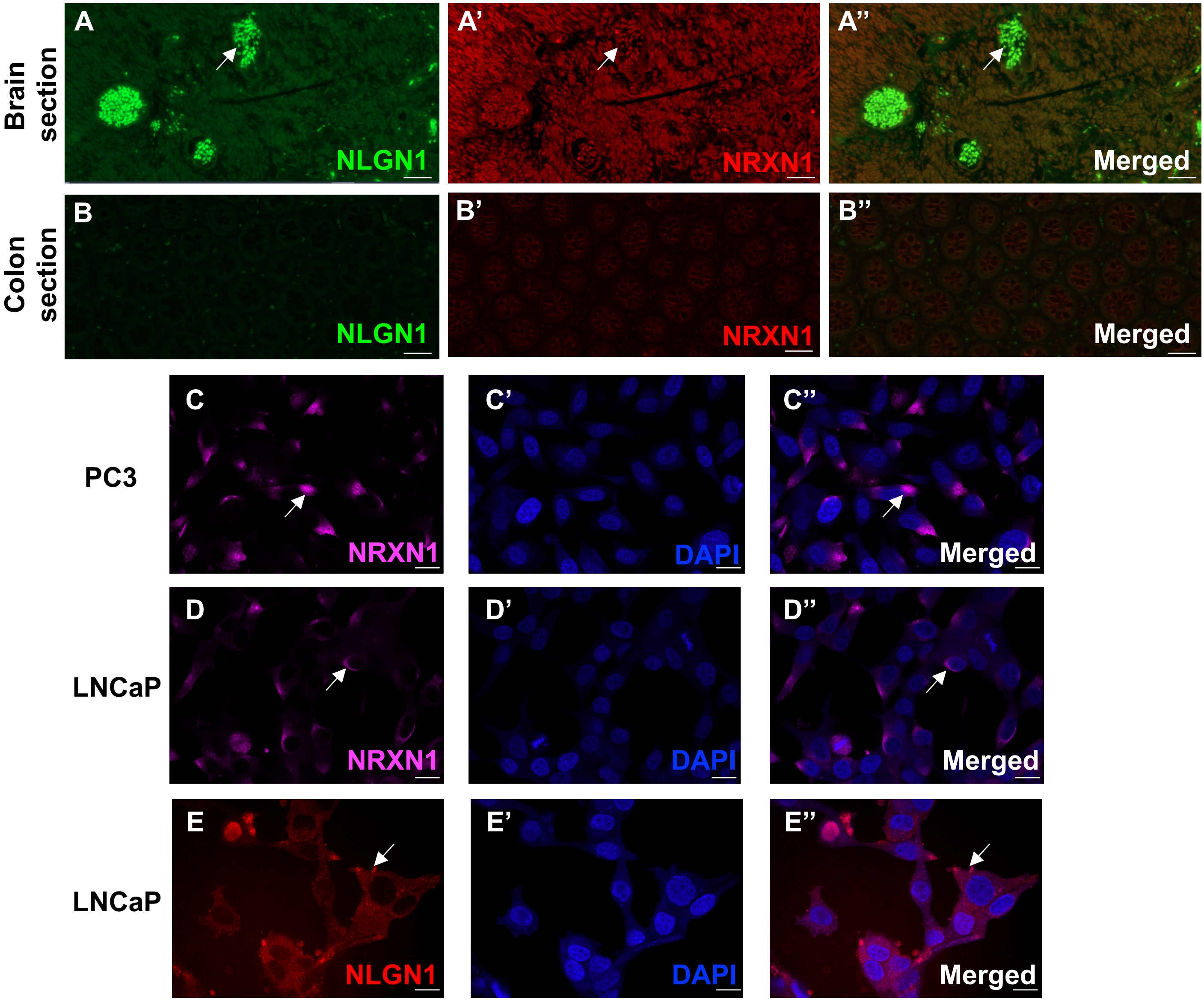


**F**

| | AF488 | AF555 | AF647 | AF750 |
|---|---|---|---|---|
| 1 | | NLGN1 | | NRXN1 |
| 2 | - | - | NCAM | CK5 |
| 3 | AF | | | |
| 4 | - | AR | CK8 | ChrA |
| 5 | - | ECAD | CD31 | CD3 |
| 6 | AF | | | |
| 7 | - | ERG | - | CK14 |

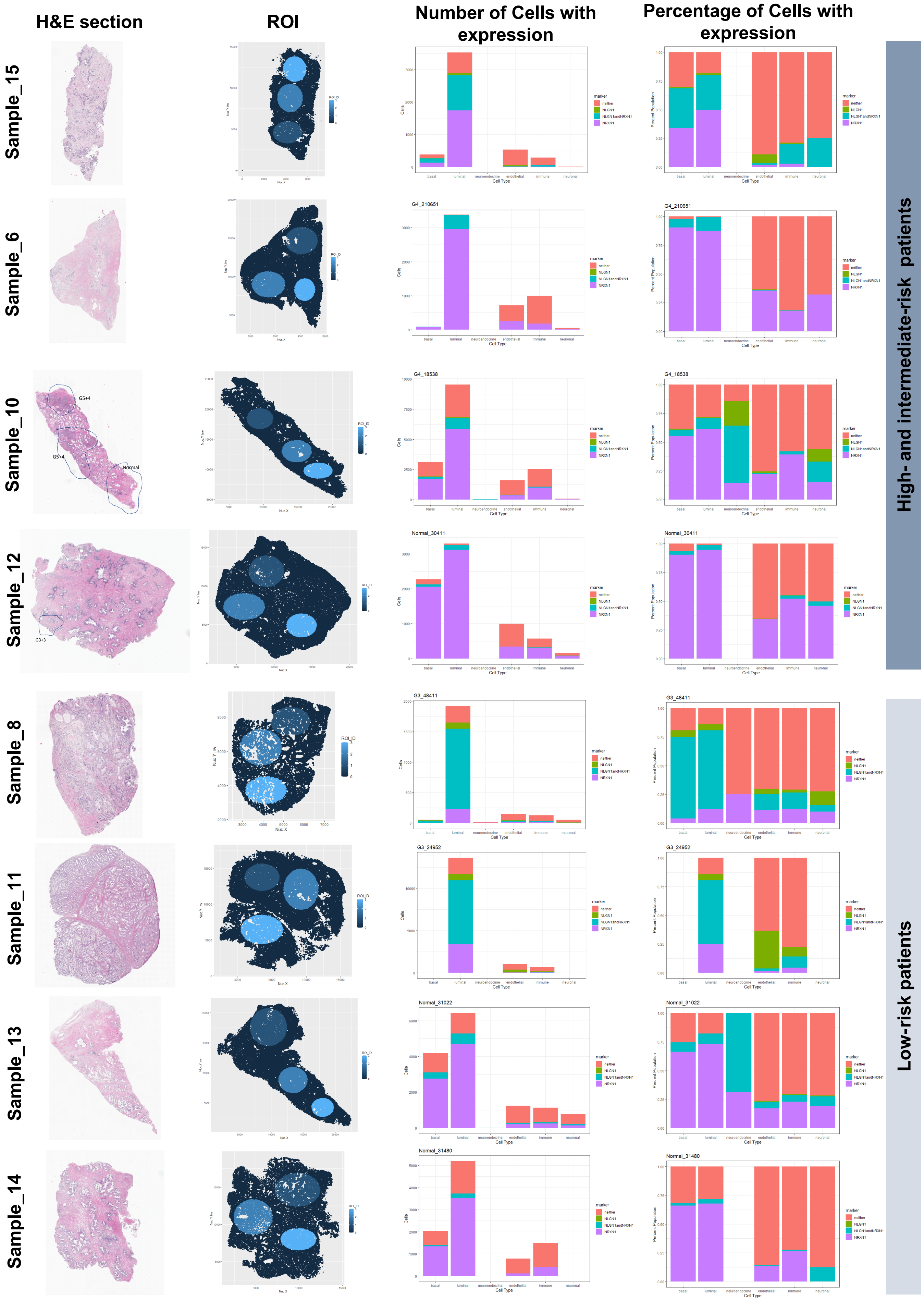
H&E section
ROI
Number of Cells with expression
Percentage of Cells with expression
Sample_15
Sample_6
Sample_10
Sample_12
Sample_8
Sample_11
Sample_13
Sample_14
High- and intermediate-risk patients
Low-risk patients
G5+4
G5+4
Normal
G3+3
G4_210651
G4_18538
Normal_30411
G3_48411
G3_24952
Normal_31022
Normal_31480
marker
neither
NLGN1
NLGN1andNRXN1
NRXN1
Cells
Percent Population
Cell Type
basal
luminal
neuroendocrine
endothelial
immune
neuronal
ROI_ID
Nuc X
Nuc Y inv
